# Pallium tracks valence-specific chemosensory amplification of eye-body coordination in larval zebrafish

**DOI:** 10.1101/2024.10.27.620486

**Authors:** Samuel K. H. Sy, Danny C. W. Chan, Jenny J. Zhang, Jing Lyu, Crystal Feng, Kui Wang, Vincent C. T. Mok, Kenneth K. Y. Wong, Yu Mu, Yu Hu, Ho Ko

## Abstract

Coordinated eye-body movements are essential for adaptive behavior, yet little is known about how multisensory input, particularly chemosensory cues, shapes this coordination. Using our enhanced Fish-on-Chips optofluidic platform, we uncovered complex dynamics in how larval zebrafish coordinate saccadic eye movements with tail flips. Under baseline conditions, spontaneous tail flips dynamically align with saccades in frequency and direction for coordinated turns. Chemosensory valence further modulates this coordination: death-associated cues intensify both the strength and relative frequency of coupled saccade-tail flips during turns, whereas food-related cues promote forward gliding without altering saccade coupling. Concurrent brain-wide neuronal imaging reveals that the zebrafish pallium represents the mapping between aversive valence and enhanced saccade-tail coordination, with stronger coupling associated with higher pallium activation. These findings uncover the neural representations and behavioral strategies by which chemosensory inputs of different valences are associated with amplified eye-body coordination and adaptive locomotion in a developing vertebrate, offering insights into principles of sensory-motor integration.

## Introduction

Eye-body coordination underpins a wide range of visuomotor behaviors across the animal kingdom, enabling animals to adapt to their environments and perform critical tasks such as hunting, navigation and foraging ^1–3^. In many vertebrates, quick and synchronous movements of both eyes in the same direction, known as conjugate saccades (hereafter referred to as saccades), play a key role in aligning gaze with body orientation, facilitating precise tracking of moving objects and stable navigation through complex environments ^4–8^. In many movement disorders, especially Parkinson’s disease, turning performance is impaired and closely linked to saccadic eye movements: turn-initiating saccades are abnormal, and their timing and kinematics explain a substantial fraction of the variance in turn duration ^9^, indicating that turning performance associates strongly with eye-body coordination, not just limb movements. Such eye-body coordination is well-illustrated in *Xenopus laevis* larvae, where compensatory eye movements during locomotion are driven by feedforward efference copies of spinal central pattern generator activity ^10^. Similar mechanisms have also been observed in mice ^11^. These signals synchronize eye movements with spinal and tail undulations to stabilize gaze during swimming while overriding reactive sensory feedback such as vestibulo-ocular reflexes ^10,12^. In humans, evidence suggests that saccades and locomotion rely on partially shared circuits spanning cortical eye fields, superior colliculus, basal ganglia, cerebellum, and brainstem premotor/locomotor centers ^5^. Although visual and mechanosensory cues are traditionally thought to dominate visuomotor control ^13–20^, an animal often needs to integrate information from multisensory modalities to optimize decision making and motor program selection ^21–25^.

Chemosensation, one of the most ancient sensory systems ^26^, plays a pivotal role in locomotion by modulating heading direction in response to chemical cues with innate and/or learnt valence values ^27–40^. These cues provide critical information about the environment, which can signal the presence of food ^28^ or predators ^29,30^, driving context-specific motor responses like fleeing or orienting. For example, in teleost fish, sudden exposure to injury-released alarm substances (“Schreckstoff”) can elicit rapid, escape-like bouts as part of a robust alarm response ^41,42^. While chemosensory valence is well-known to shape body movements, its influence on eye movements and eye-body coordination remains poorly understood. In addition, chemical cues may trigger arousal-related state changes, potentially leading to more frequent visual scanning and heightened alertness ^43^. Understanding how chemosensory cues interact with visuomotor behaviors could provide key insights into the mechanism of multisensory integration and motor control.

In this study, we investigated how chemosensory valence modulates the coordination of eye and body movements in larval zebrafish. Zebrafish are an ideal model for studying sensorimotor integration due to their lack of a flexible neck (i.e., the anatomy limits independent yaw (side-to-side) movement of the head), which simplifies kinematic analysis ^44,45^. Their optical transparency also allows for brain-wide neuronal imaging, making them highly suitable for studies of neural circuit activity ^46^. Beyond these technical advantages, larval zebrafish naturally use waterborne chemosensory cues to guide turning and forward swimming. Vision is widely believed to have first evolved in aquatic environments ^47^, and dissolved chemicals and light gradients jointly shaped early sensorimotor control, making waterborne cues a natural context for studying their integration. Although odor dispersal in water differs from that in air, animals in both media must integrate changing chemical signals with visual and mechanosensory inputs to select appropriate locomotor responses. We hypothesized that chemosensory cues trigger a coordinated behavioral response in larval zebrafish with saccades and tail flips jointly modulated by cue valence. To test this, we advanced our Fish-on-Chips system ^27^, enabling simultaneous behavioral analysis and brain-wide imaging and efficient chemical valence screening in freely responding larvae. This approach allowed us to elucidate the neural mechanisms by which chemosensory input regulates coordinated visuomotor behaviors.

We revealed that temporally coupled saccade-body turns are characterized by a larger turn intent, and exhibit dynamic directional alignment and temporal coordination. In contrast, these characteristics are not observed in saccade-independent forward swims. Upon exposure to chemosensory stimuli, coupled saccade-turn is enhanced by aversive death-associated but not appetitive food-related chemicals. Across the brain, we identified neuronal signatures underlying chemosensory input-amplified coupled saccade-tail flips, suggesting the telencephalic pallium as a candidate region. Our findings indicate that in larval zebrafish, chemosensory cues can influence eye-body coordination based on innate perceived valence, serving as an integrated response to achieve locomotive goals.

## Results

### Fish-on-Chips 2.0 enables efficient valence screening and naturalistic behavior-neural imaging under chemical cue delivery

Despite tremendous efforts in developing appropriate methodology for the study of chemosensation in small animals, previous methods ^48–51^, including our own ^27^, have fallen short of achieving high-throughput, spatiotemporally precise valence screening and naturalistic behavioral readout under repeated delivery of multiple chemical cues—a challenge that we address in our study. To address this gap, we advance our Fish-on-Chips platform to demonstrate significantly improved experimental conditions for the mechanistic studies of behavioral and neural effects of chemicals on larval zebrafish.

First, we developed an improved valence assay (see **Methods, Figures 1a, b, Supplementary Figure 1a**) that allowed us to efficiently screen preferences of ecologically significant chemicals or odors across multiple orders of concentrations using the same animals, covering wide dynamic ranges of chemoreception in teleosts ^52^. These included two amino acids, namely valine (Val) and alanine (Ala) ^32,33^, the sex pheromone prostaglandin F2α (PGF2α) ^31^, putrefaction-associated diamines including cadaverine (Cad) and putrescine (Put) ^27,29^, bile acids ^35,36^ including chenodeoxycholic acid (CDCA) and glycodeoxycholic acid (GDCA), adenosine (Ade) which is a food-associated cue ^28^, and sodium chloride (NaCl) which determines salinity ^34^.

**Figure 1.**
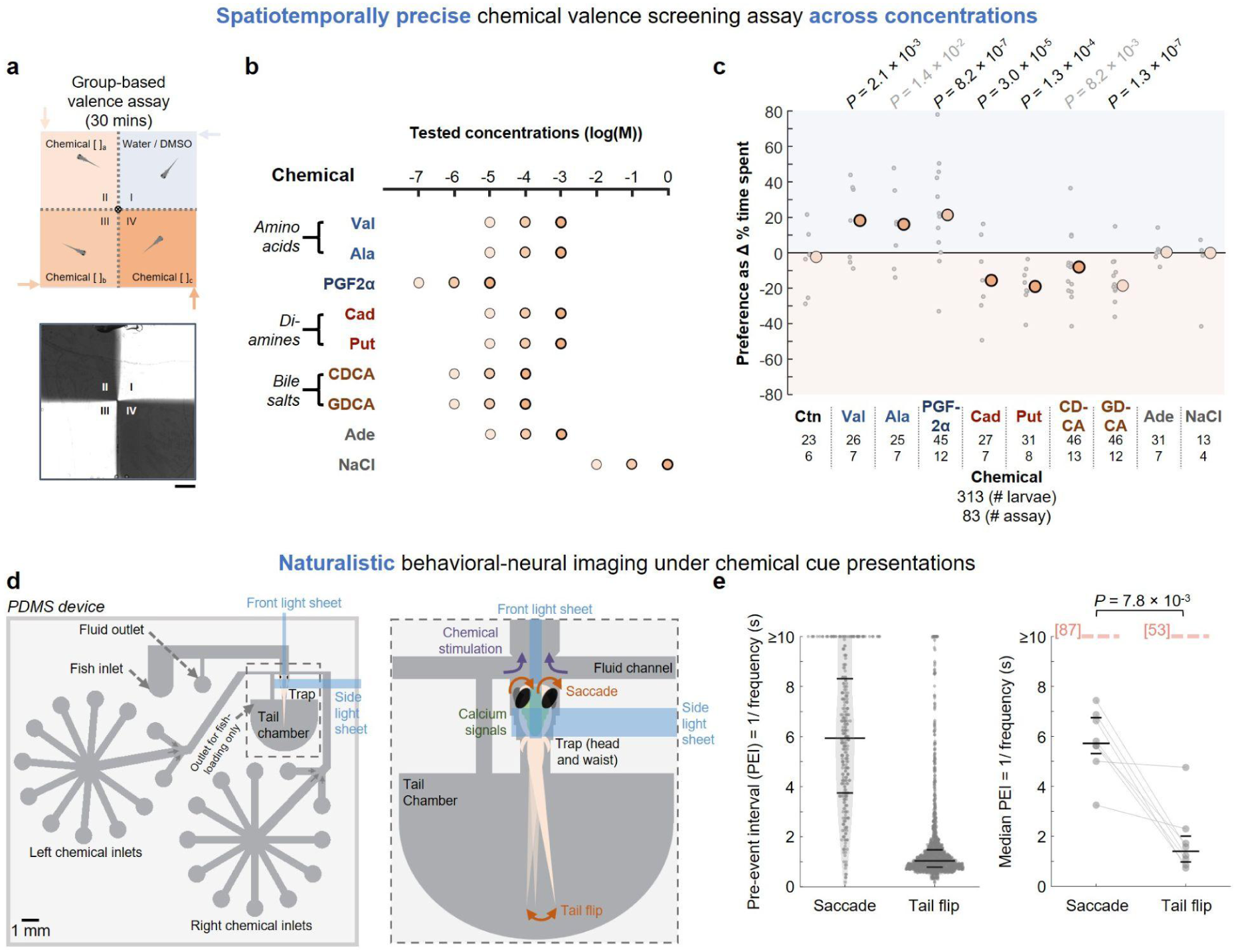
Fish-on-Chips 2.0 for studying chemosensation in larval zebrafish. **(a)** Upper panel: Schematic of the valence assay. Zebrafish larvae swimming in a two-dimensional arena (40 mm × 40 mm × 1.5 mm) are imaged at 20 fps under infrared (IR) illumination in the absence of visible light (see **Supplementary Figure 1a** for a more detailed schematic of the setup). Four quadrants with equal area are created and maintained by a constant, slow inflow of fluid at each corner (marked by arrows), with outflow occurring at a shared central outlet. The laminar flow maintains static borders between the zones. Lower panel: Visualization of the well-defined quadrants by infusing IR dyes into quadrants II and IV, and water into quadrants I and III. Scale bar: 5 mm. **(b)** Assayed chemicals and tested concentrations (varied across 3 consecutive orders of magnitude for each), including valine (Val), alanine (Ala), prostaglandin F2α (PGF2α), cadaverine (Cad), putrescine (Put), chenodeoxycholic acid (CDCA), glycodeoxycholic acid (GDCA), adenosine (Ade), and sodium chloride (NaCl). These were dissolved either in water or 0.5% DMSO and infused into quadrants II (lowest concentration) to IV (highest concentration). **(c)** Differences in % time spent in the most representative chemical quadrant (concentration indicated by the dot color) compared with the control quadrant I, showing the values of individual assays (gray dots) and the medians (black-outlined dots). Results of all quadrants are shown in **Supplementary Figure 1c**. Below each chemical name, the first number indicates the total number of larvae assayed, and the second number specifies the total number of assays performed. *P*-values: For each chemical, multiple *Z*-tests comparing mean differences in % time between the control quadrant I and all individual non-control quadrants and their possible combinations, including the representative quadrant shown, against a null hypothesis of zero difference. The displayed *P* value is from the most significant comparison involving that representative quadrant. The population standard deviation (S.D.) was estimated from the sample S.D. of the control group (see **Methods**). Statistical significance was determined using a Benjamini–Hochberg-derived significance threshold applied to the unadjusted *P*-values (controlled within each odor). Black, significant; gray, near-significant. **(d)** Left panel: A drawing of the enhanced polydimethylsiloxane (PDMS)-based microfluidic device, featuring multiple chemical delivery channels on both sides of the larval head chamber. These channels are offset from the front to accommodate front excitation light sheet scanning, in addition to the right side light sheet. Right panel: A close-up illustration of the neural-behavioral imaging capabilities in a larval subject during precisely controlled chemical stimulus presentation. See **Supplementary Figure 2** for more details. **(e)** Left panel: Pre-event intervals (PEIs) [1/ frequency] of spontaneous saccade and tail flip for all recorded events in the microfluidic device. Shaded areas scale according to the probability density function of values. Data collected from *n* = 8 larvae with 296 spontaneous saccades and 1,509 spontaneous tail flips, each with a recorded preceding reference event. Right panel: Median pre-event intervals (PEIs) [1/ frequency] of spontaneous saccade and tail flip of each larva. Gray lines connect the paired saccade and tail-flip median PEIs from the same larva. *P*-value: Two-sided Wilcoxon signed-rank test on paired larva-level median PEIs (*n* = 8 larvae). Dotted rose lines indicate the reduced frequencies typically reported in studies using agarose-based embedding, both < 0.1 Hz (see refs ^87^ and ^53^ for saccades and tail flips respectively). In both panels, horizontal lines show the median, 25^th^, and 75^th^ percentiles for each group.

In control conditions with all quadrants filled with water, the larval zebrafish distribute their time evenly across the four quadrants, indicating no intrinsic bias in the setup (**Figure 1c**, **Supplementary Figure 1c**). We observed that Val (at 100 μM or above) and Ala (at 10 μM and 1 mM) are attractive to the larvae, similar to the preference seen in adult zebrafish ^32,33^. Interestingly, despite being a sex pheromone ^31^, PGF2α is already attractive to the larval zebrafish at a concentration of 1 μM. We also confirmed that Cad and Put, at 10 μM or 100 μM and above, respectively, are repellent ^27,29^. CDCA (at 10 μM and above) and GDCA (at 1 μM and above), which are examples of bile acids typically considered as social cues at low concentrations ^35,36^, were found to be repellent. For PGF2α and the bile acids, which require 0.5% DMSO for dissolution, we obtained consistent results comparing data using either water or 0.5% DMSO in the control quadrant (**Supplementary Figures 1d, e**), and therefore pooled the results for analysis (**Figure 1c, Supplementary Figure 1c**).

Notably, larval zebrafish generally respond to attractive and aversive chemicals at the highest concentration tested, with the exception of PGF2α. Adenosine, known to attract adult zebrafish ^28^, did not exhibit any attractant or repellent effects on the larvae. Furthermore, NaCl, a recently reported aversive olfactory stimulus ^34^, showed no impacts on the larvae in our assays. Note that however, our assay predominantly presents absolute concentrations of NaCl with minimal gradient, which could be a critical factor as zebrafish may rely on its spatial gradient for navigation to avoid high salinity areas ^34^.

Next, in our new imaging setup for enhanced larva tethering, chemical delivery and brainwide neuronal imaging abilities (see **Methods, Figure 1d, Supplementary Figure 2**), we observed robust spontaneous saccades and tail flips (during blank trials and pre-stimulus windows) across 8 larval subjects (**Figure 1e**), with tail flip frequency comparable to that of freely moving larvae ^53^. Notably, spontaneous saccades occurred less frequently than spontaneous tail flips, as indicated by the longer median pre-event interval (PEI, defined as difference in onset times between an event and the preceding one) for saccades (5.94 seconds; *n* = 296 saccades from 8 larvae) vs. tail flips (1.04 seconds; *n* = 1,509 tail flips from 8 larvae). Since these behaviors preserve more naturalistic behavior (in terms of saccade and tail flip frequencies) compared to that reported with the conventional agarose-based tethering approaches (**Figure 1e**), the setup is well-posed for the study of both the behavioral and neural effects under highly controlled chemical stimulus delivery.

### Tail flip dynamically aligns with saccade in rate and direction for spontaneous turns

As mentioned, numerous studies have shown that eye and body movements are precisely coordinated during single spontaneous orientation changes in larval zebrafish ^6,7^ and other vertebrates. However, it remains unclear how saccades and tail flips—two quasi-periodic movements that occur at vastly different rates (**Figure 1e**)—are coordinated during self-initiated turns. Interestingly, this coordination contrasts with predictions of smooth and pre-planned movements in some motor control models ^54,55^. We hypothesize that this coordination extends beyond individual events and requires the dynamic spatiotemporal matching of the more frequently occurring tail flips to saccades. To test this, we analyzed the recorded baseline behaviors.

We first isolated the co-occurrences of saccades and tail flips, defining temporally coupled saccade-tail flip events (hereafter referred to as S-T events) as those with saccade and tail flip onset time difference of 0.5 seconds or less. This threshold is based on it being approximately half the median PEI of the more frequent tail flips. Other non-coupled saccade or tail flip events are referred to as independent events. We first examined the relative timings of saccade and tail flip in an S-T event (**Figure 2a**). The tail flip typically occurred after the coupled saccade (**Figure 2b**), with their durations largely overlapping (**Figure 2c**), indicating that they represent coordinated eye-body movements at individual event levels. Overall, 52.8% of saccades and 15.0% of tail flips occurred in S-T events (**Supplementary Figure 3a**). Beyond single events, there is evident temporal coordination in their rate compared with when saccades and tail flips occur independently, for which these intervals remained relatively stable (**Figures 2d, e**). Following an S-T event, the PEI for the subsequent saccade increased by ∼23% (**Figure 2d**), effectively extending the time for gaze fixation. For tail flips, their PEI around S-T events significantly lengthened, increasing by ∼50% in magnitude from 6 prior tail flips to the peak at S-T events, then decreasing back to the baseline over a few tail flips (**Figure 2e**). This pattern indicates that temporal control in S-T events is based on anticipatory and dynamic regulation extending beyond individual occurrence, similar to many other timed behaviors ^56^.

**Figure 2.**
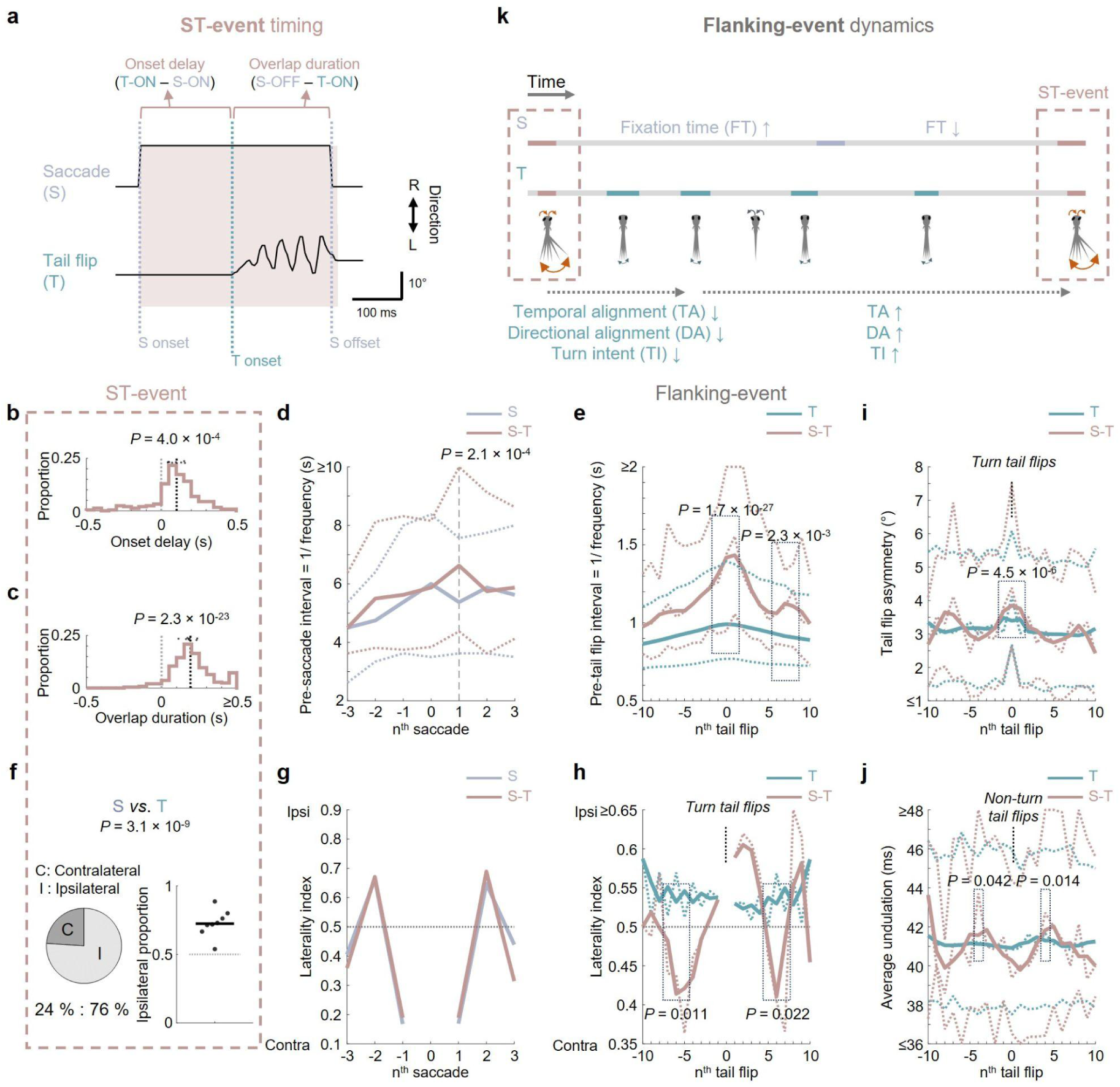
Tail flip dynamically aligns with saccade in rate and direction for performing spontaneous body turns. **(a)** Schematic illustrating the temporal parameters of an example coupled saccade-tail flip (S-T) event. **(b)** & **(c)** Histograms of **(b)** onset delay and **(c)** overlap duration for coupled saccade-tail flip (S-T) events. Note that the precision of measurements for the onset and offset for saccade and tail flip were 0.0625s and 0.0025s due to the respective sampling rates. Black and gray dashed lines indicate medians and zero, respectively. Dots indicate the median delay for each larva. *P*-values: Linear mixed-effects model with a random intercept per larva, testing the fixed intercept against zero. **(d)** & **(e)** Pre-event intervals [1/ frequency] of self and flanking **(d)** saccades and **(e)** tail flips with respect to independent (S, saccade; T, tail flip) or S-T events. **(f)** Left panel: Pie chart showing the proportions of contra- and ipsilateral turns, comparing the directionality of saccades (S) and tail flips (T; only turns considered, see **Methods**) in S-T events. Right panel: Larva-based ipsilateral proportion. The horizontal line indicates the median across larvae. *P*-value: Binomial generalized linear mixed-effects model with a random intercept per larva (ipsilateral vs. contralateral), testing whether the ipsilateral fraction differs from 0.5. **(g)** & **(h)** Laterality indices (defined as proportions of ipsilateral turns) in flanking **(g)** saccades and **(h)** turn tail flips (tail flip asymmetry ≥ 1.5°) with respect to independent (S, T) or S-T events. In **(h)**, the raw proportions (thin dotted lines) and corresponding moving-average curves (thick solid lines) are shown. *P*-values: For the peak lag and its adjacent ±1 lags, binomial generalized linear mixed-effects models with a random intercept per larva (ipsilateral vs. contralateral), testing whether the ipsilateral fraction differs between S-T-associated and independent events. **(i)** Tail flip asymmetry of self and flanking tail flips for independent tail flips (T) or S-T events with tail flip asymmetry ≥ 1.5° (see **Supplementary Figure 3e** for all events). **(j)** Average undulation of self and flanking tail flips for independent tail flips (T) or S-T events with tail flip asymmetry < 1.5° (see **Supplementary Figure 3f** for all events). **(k)** Conceptual summary of the temporal, directional, and kinematic coordination between spontaneous saccades and tail flips during body turns. In coupled saccade–tail-flip (S-T) events, saccades begin shortly before tail flips, such that the two movements largely overlap in time. Tail flips preferentially turn to the same side as the saccade and exhibit stronger turn-like kinematics. Across movements surrounding an S-T event, the temporal coordination of event rates, directional alignment, and tail-turn kinematics progressively strengthen toward the S-T event, peak at the event, and weaken thereafter. Following an S-T event, the interval until subsequent saccades is prolonged, consistent with extended gaze fixation. Abbreviations in **(a)**, ON, onset; OFF, offset. In (**d)**, the two thin dotted lines represent the 25^th^, and 75^th^ percentiles of all events, whereas the thick solid lines represent the medians. In **(e)**, **(i)**, and **(j)**, the thin dotted lines represent the medians, 25^th^, and 75^th^ percentiles of all events, and the thick solid lines show the corresponding moving-average curves of the medians. *P*-values: Linear mixed-effects models with event type as a fixed effect and larval identity as a random intercept, applied to events pooled across the peak lag and/or its adjacent ±1 lags. For all plots, data were collected from *n* = 8 larvae, with 472 spontaneous saccades and 1,663 spontaneous tail flips.

During single S-T events, saccades and tail flips are predominantly ipsilateral (i.e., towards the same body side) (**Figure 2f**), in line with previous reports ^6,7^. Considering the dynamic nature of saccade-tail flip coordination, we investigated the directional synchronization of these movements, by analyzing the directionality of both movements in independent and coordinated events. Consecutive saccades mostly alternated in direction (**Supplementary Figure 3b**), and such a pattern did not differ between independent saccades vs. those of S-T events (**Figure 2g**). In contrast, tail flips generally exhibited an ipsilateral tendency relative to each preceding flip (**Supplementary Figure 3c**) and maintained this pattern across a series of tail flips (**Figure 2h, *teal curve***). This aligns with previous reports of history-dependent tail-flip direction ^57^ and stable individual left-right motor asymmetries in larval zebrafish ^58^. Interestingly, when aligned to S-T events, the dominant direction of tail flip gradually alternated from and to the contralateral side over ±5–6 tail flips (**Figure 2h, *coral curve***). As the median pre-saccade interval was also around 6 times that of pre-tail flip interval (**Figure 1e**), this gradual modulation likely permits directional alignment of tail flips with consecutive saccades.

We further analyzed the relationship between S-T events and putative locomotive intent using kinematic parameters of each tail flip extracted from time-series tail angle data (**Supplementary Figure 3d**). Tail flip asymmetry gradually increased before and peaked at S-T events, followed by a decrease over a few tail flips (**Figure 2i, Supplementary Figure 3e**). This pattern appeared to recur approximately every 6–7 tail flips (**Figure 2i, Supplementary Figure 3e**). Changes in the average undulation exhibited an antiphasic pattern to that of tail flip asymmetry, peaking at around ±3^rd^–4^th^ tail flips relative to S-T events (**Figure 2j, Supplementary Figure 3f**). Together, these indicated that S-T events are associated with a greater orientation towards turning (see also similar/related findings in ^6,7^) rather than sustained forward gliding ^59^, with the kinematics of tail flips modulated in an anticipatory manner along with S-T event timing.

Collectively, we concluded that in larval zebrafish, spontaneous coupled eye-tail flip movements involve highly organized motor coordination. This coordination, occurring through a sequence of tail events, is characterized by simultaneous occurrence of temporal synchronization, directional alignment, and kinematic modulation (**Figure 2k**), putatively for mediating body turns during locomotion.

Apart from saccade-coupling, the above-mentioned findings implicated that spontaneous swim bouts are dynamically organized around turns. Using the swim bout data alone from our previous study ^27^, we classified the swim bouts into non-turn and turn bouts (see **Methods**), as surrogates for independent and saccade-coupled tail flips, respectively (recall that saccade-coupled tail flips have larger turning intent, **Figure 2i, Supplementary Figure 3e**). Upon comparative analysis, we found similar temporal (**Supplementary Figure 4a**), directional (**Supplementary Figure 4b**) and kinematic (**Supplementary Figure 4c**) coordinations for the turn bouts, but not the non-turn bouts (**Supplementary Figure 4**). The behavioral manifestations of larval zebrafish in a freely moving arena therefore further substantiated that observed in the microfluidic device.

### Chemical valence differentially modulates saccade-tail flip coordination

We next aimed to test whether chemical stimuli, often encountered unexpectedly in the natural environment, may influence eye movements and saccade-turn coordination. In response to aversive chemical stimuli, turns may be co-modulated with saccades, and this could in theory manifest in several different yet mutually non-exclusive ways. For example, as S-T events likely mediate turns, the proportion of saccade-coupled tail flips can be increased. In addition, kinematic coordination during S-T events can be enhanced to make more pronounced turns. On the other hand, turns and forward gliding can also be modulated independent of S-T coordination, by altering the overall tail flip asymmetry and undulation interval of tail flips. To test these possibilities, we used potent and ethologically representative appetitive or aversive chemical stimuli determined by the previous valence assays, including the death-associated diamines (Cad and Put) and the food-related amino acids (Val and Ala), each at a concentration of 1 mM (**Figure 1c, Supplementary Figure 1c**). Each of these chemicals and a blank control were precisely administered to larval zebrafish for a duration of 10 seconds in a pseudorandomized order with resting intervals during neural-behavioral imaging sessions (**Figure 3a**, see **Methods**).

**Figure 3.**
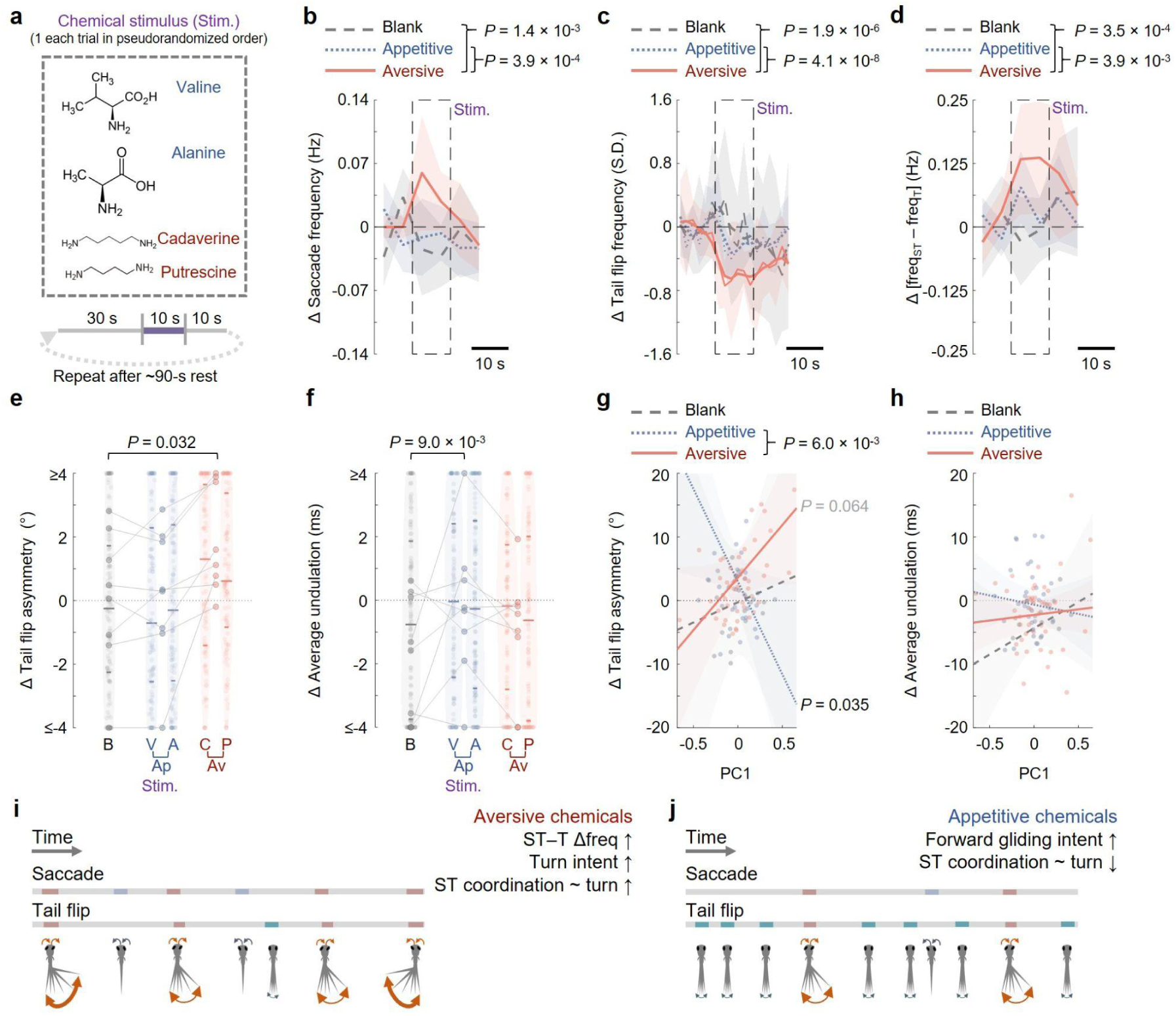
Aversive but not appetitive chemosensory cues amplify eye-body coordination via modulations of individual and joint movements of saccades and tail flips. **(a)** Illustration of chemical stimulation paradigm. During each trial, one appetitive or aversive chemical cue, or a blank control, was administered to a tethered larval subject under neural-behavioral imaging. (b)–(d) Stimulus-evoked changes in (b) saccade frequency, (c) normalized tail flip frequency (*z*-score relative to the pre-stimulus baseline; see Methods), and (d) relative frequency of coupled saccade-tail flip (S-T) events vs. independent tail flips (i.e., Δ[S-T event frequency - independent tail flip frequency]) across time, compared with the pre-stimulus average. (e) & (f) Stimulus-evoked changes in (e) tail flip asymmetry (all tail flips) and (f) average undulation (for tail flips with ≥ 4 undulation cycles), also relative to baseline. Analyses were performed on tail flips occurring during the 10-s stimulus window. (g,h) (g) Δ tail-flip asymmetry and (h) Δ average undulation (for S-T events whose tail flips had ≥ 4 undulation cycles) plotted against PC1 score of S-T temporal coordination for S-T events occurring during the 10-s stimulus window. Lines show mixed-effects model fits per group and shadows show 95% confidence intervals. PC1 was derived from five temporal coordination variables (see Methods and Supplementary Figure 6). Only events with saccade onset certainly earlier than tail-flip onset were included (see Methods). (i) & (j) Illustrations of larval zebrafish behavioral changes upon (i) aversive and (j) appetitive chemical presentations. In (b)–(d), quantities are binned and normalized to the average values in the 10-second pre-stimulus windows (see Methods). Lines show means and shaded bands show 95% confidence intervals across larvae; for each larva, values were averaged across repeated trials within each stimulus condition. In (c), the raw means (thin lines) and corresponding moving-average curves (thick lines) are shown. Dashed rectangles mark the stimulus window. In (e) & (f), horizontal lines indicate medians, 25^th^, and 75^th^ percentiles. Shadows of the violin plots scale according to the probability density function. Larger dots indicate the median for each larva in each condition, and gray lines connect the same larva across stimulus conditions, illustrating within-animal changes. In (b)–(f), *P*-values: linear mixed-effects models with group as a fixed effect and larvae as a random effect. For (b)–(d), statistical tests were performed on data from the stimulus window and the subsequent 5 s post-stimulus period (see Methods). In (g) & (h), lines and confidence intervals are from linear mixed-effects models of Δ metrics versus PC1 with PC1, PC2, group, and their interactions with group as fixed effects and larvae as a random effect. *P*-values: PC1 slopes against zero within groups and differences in PC1 slopes between appetitive and aversive conditions. Abbreviations: Ap, appetitive; Av, aversive; B, blank; V, valine; A, alanine; C, cadaverine; P, putrescine. For all plots, data were collected from *n* = 8 larvae, and the total numbers of blank control, appetitive and aversive chemical trials used in statistical analyses are 45, 83 and 86, respectively. In (c), the numbers of blank, appetitive and aversive trials with non-zero pre-stimulus tail flip frequency used in statistical analyses are 28, 52 and 46, respectively. In (e), the numbers of blank, appetitive, and aversive events analyzed were 141, 244, and 214, respectively. In (f), the corresponding numbers were 127, 227, and 196.

We first examined the effects of chemical stimuli on saccade, putative turn and forward motion individually. Only the aversive chemicals led to a higher saccade frequency and a lower tail flip frequency during their presentation (**Figures 3b, c**), both of which returned to baseline levels shortly after the cessation of stimulation (**Figures 3b, c**). During aversive stimuli, the proportion of saccade-coupled tail flips increased (**Figure 3d**), along with an increase in tail flip asymmetry (**Figure 3e**). In contrast, appetitive stimuli did not lead to any discernible change in tail flip asymmetry (**Figure 3e**). To examine whether larvae may intend to perform more forward swims in response to any chemical stimulus, we analyzed tail flips with ≥ 4 undulation cycles. We observed that only appetitive stimuli significantly increased the average undulation of these tail flips (**Figure 3f**), presumably leading to more sustained forward gliding movements ^59^. Overall, although the individual effects on saccades, tail flips, and stimulus trials appeared modest (**Figures 3b–f**), similar small, context-dependent biases in locomotor behavior have been shown to accumulate over time to produce robust navigational outcomes, as demonstrated in thermosensory navigation in larval zebrafish.^60,61^ Thus, even subtle shifts in saccade frequency, coupling, and tail kinematics could cumulatively bias behavior toward avoidance or approach when integrated across successive bouts.

In the presence of chemical stimuli, saccades and tail flips during S-T events showed broadly similar temporal (**Supplementary Figures 5a–d**) and directional (**Supplementary Figures 5e, f**) alignments. Saccade-coupled tail flips also exhibited generally greater tail flip asymmetry and smaller average undulation than the independent tail flips (**Supplementary Figures 5g, h**), consistent with the turning-associated kinematics of S-T events.

Because aversive stimuli increased saccade occurrence (**Figure 3b**), we tested whether the structured features of S-T events could arise through chance temporal co-occurrence. We jointly circularly shifted saccade onset and offset times within individual trials, preserving saccade count, duration, direction, and the tail-flip sequence, before reidentifying S-T events using the original criterion. The shuffled aversive-trial S-T events did not reproduce the saccade-leading timing or ipsilateral eye–tail alignment of observed S-T events (**Supplementary Figures 5i, j**). Together, these results indicated that altered event rates alone are unlikely to fully explain the temporal and directional organization of aversive-context S-T events.

Next, we asked whether temporal features of S-T events, including saccade and tail-flip onset and offset delays, their respective durations, and the S-T overlap duration, are differentially regulated as a function of chemosensory valence to adjust turn magnitude. Because these temporal parameters were highly correlated (**Supplementary Figure 6a**), we applied principal component analysis (PCA) to this set of variables and used the first two principal components (PC1 and PC2) as predictors in a linear mixed-effects model. PC1 explained 61% of the variance and was dominated by opposite loadings of S-T onset/offset delays and overlap duration, such that shorter onset/offset delays and greater overlap yielded higher PC1 scores, whereas PC2 captured a smaller (24%) but orthogonal component of the temporal structure (**Supplementary Figures 6b, c**). Thus, PC1 primarily reflects the temporal proximity of saccades and tail flips. Chemosensory valence bidirectionally modulated the relationship between PC1 and chemical-evoked change in tail flip asymmetry: aversive stimuli produced a positive relationship, whereas appetitive stimuli produced a negative one (**Figure 3g**). These valence-dependent effects were not observed for chemical-evoked change in average undulation for S-T events whose tail flips had ≥ 4 undulation cycles (**Figure 3h**), suggesting that S-T temporal proximity is specifically regulated to adjust turn magnitude in response to aversive cues.

Taken together, the two opposite chemical valences elicit distinct coordinated behavioral responses in larval zebrafish. Aversive chemicals lead to an increase in saccades and a higher proportion of enhanced, temporally coordinated saccade-coupled tail flips (**Figure 3i**), whereas appetitive chemicals result in more effective forward gliding movements without changes in saccade patterns or tail flip coupling (**Figure 3j**). These findings highlight the flexibility of the eye-body coordination system in zebrafish larvae, adapting to different chemical cues.

### Pallial neuronal subsets are highly co-tuned to S-T events and chemical aversiveness

By analyzing the concurrently recorded brain-wide neuronal activities in 5 out of 8 behaviorally recorded larval subjects (**Figures 4a, b**), we aimed to identify neuronal ensembles involved in the control of coupled saccade-tail flips. To this end, we first designed motor regressors for each larva (see **Methods**). By calculating the mutual information (M.I.) between neuronal activity and these regressors, we identified neurons encoding spontaneous tail flips, and determined their preferential involvement in independent vs. saccade-coupled tail flips (see **Methods** for definition of motor preference). This analysis revealed that motor neurons across the brain generally exhibited a preference for independent tail flips, with a notable concentration in key motor command centers, specifically the brainstem mesencephalon and rhombencephalon ^62,63^ (**Figures 4c, d**). Interestingly, neurons in the telencephalic pallium, an evolutionarily conserved dorsal telencephalic structure that gives rise to cortical, hippocampal, and amygdala-like circuits in more complex vertebrates ^64^, displayed the weakest preference for independent tail flips and the strongest relative preference for S-T events, compared to other brain regions (**Figures 4c, d**).

**Figure 4.**
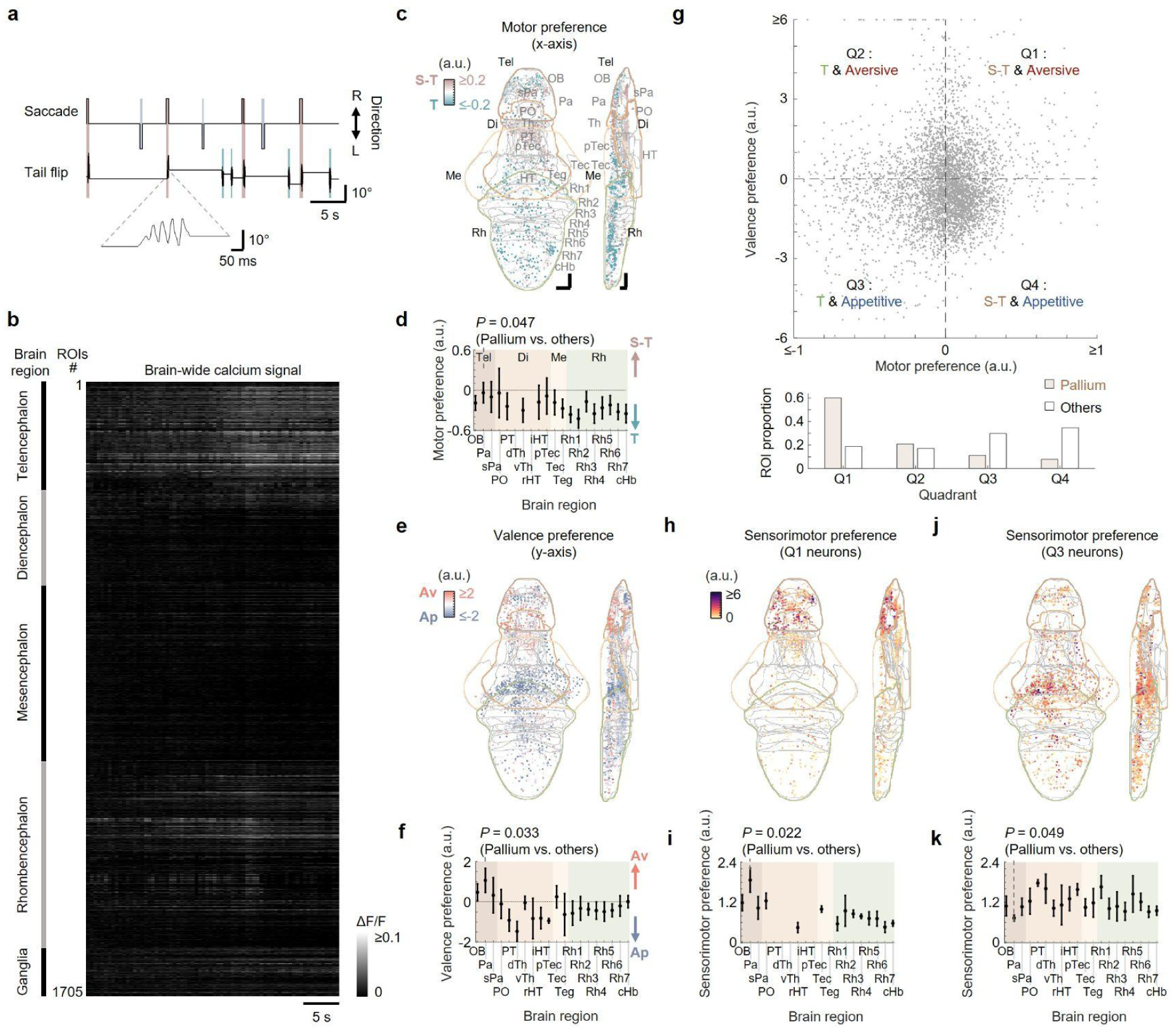
Pallial neuronal subsets are highly co-tuned to coupled saccade-tail flip (S-T) events and chemical aversiveness. **(a)** Spontaneous saccades and tail flips of a larval subject, with a zoom-in example tail flip shown below. Temporally coupled saccade-tail flips (≤ 0.5s onset time difference) are indicated with coral blocks. Other saccades and tail flips are highlighted with blue and teal blocks, respectively. **(b)** Heatmap of brain-wide spontaneous neuronal activities of the same larval subject in **(a)**, acquired simultaneously with the motor outputs shown. **(c)** Mean intensity projections (to transverse and sagittal planes) of motor preference of motor-encoding regions-of-interest (ROIs) (see **Methods**, x-axis of the upper panel in **(g)**). **(d)** Regional mean ± S.E.M. (across larvae) of motor preference, averaged across ROIs within each region and larvae. **(e)** Mean intensity projections (to transverse and sagittal planes) of valence preference for all valence-encoding ROIs (see **Methods,** y-axis of the upper panel in **(g)**). **(f)** Regional mean ± S.E.M. (across larvae) of valence preference, averaged across ROIs within each region and larvae. **(g)** Upper panel: Scatter plot of valence preference vs. motor preference of the union set of individual motor- and valence-encoding neurons. The four quadrants classify neurons into distinct sensorimotor tuning properties. Lower panel: Proportions of ROIs in each sensorimotor quadrant for pallial and non-pallial brain regions. ROI counts for quadrants I–IV are 107/37/20/14 (pallium) and 798/730/1276/1484 (non-pallial regions). Data are pooled across *n* = 5 larvae. **(h)** Mean intensity projections (to transverse and sagittal planes) of norm of sensorimotor preference (see **Methods**), for all ROIs preferring S-T event and aversive valence (i.e., quadrant I in **(g)**, also see **Supplementary Figure 8c** for zoom-in of the forebrain area). **(i)** Regional mean ± S.E.M. (across larvae) of norm of sensorimotor preference for all ROIs preferring S-T event and aversive valence, averaged across ROIs within each region and larvae. **(j)** Mean intensity projections (to transverse and sagittal planes) of the norm of sensorimotor preference for all ROIs preferring independent tail flip and appetitive valence (quadrant III in **(g)**, also see **Supplementary Figure 8d** for zoom-in of the forebrain area). **(k)** Regional mean ± S.E.M. (across larvae) of the norm of sensorimotor preference, for all ROIs preferring independent tail flip and appetitive valence, averaged across ROIs within each region and larvae. Scale bars in **(c)**: 50 μm in the Z-Brain Atlas space. In **(d)**, **(f)**, and **(i)**, *P*-values: Right-sided paired *t*-test comparing the pallium and non-pallial brain regions at the larval level (*n* = 5 larvae). In **(k)**, *P*-value: Left-sided paired *t*-test comparing the pallium and non-pallial brain regions at the larval level (*n* = 5 larvae). In **(c)**–**(f)** & **(h)**–**(k)**, the major brain regions (Tel: telencephalon, Di: diencephalon, Me: mesencephalon and Rh: rhombencephalon) are color-coded. Abbreviations of brain regions: OB olfactory bulb, Pa pallium, sPa subpallium, PO preoptic area, PT posterior tuberculum, dTh dorsal thalamus, vTh ventral thalamus, rHT rostral hypothalamus, iHT intermediate hypothalamus, pTec pretectum, Teg tegmentum, Rh1 rhombomere 1, Rh2 rhombomere 2, Rh3 rhombomere 3, Rh4 rhombomere 4, Rh5 rhombomere 5, Rh6 rhombomere 6, Rh7 rhombomere 7, cHb caudal hindbrain. Other abbreviations: S-T, coupled saccade-tail flip; T, independent tail flip; Ap, appetitive; Av, aversive. For all plots, neural activity data are from *n* = 5 larvae and correspond to a subset of the behavioral dataset shown in Figures 2 & **3** (*n* = 8).

We then examined the spatial distributions of chemical valence-encoding neurons (**Supplementary Figure 7**) (see **Methods** for definition of valence preference). Among the primary projection target regions of the olfactory bulbs in the telencephalon ^65,66^, we observed distinct activation patterns in response to aversive and appetitive chemicals (**Figures 4e, f, Supplementary Figure 7**). Aversive chemical-elicited activities are channeled through both the pallium and subpallium, whereas appetitive chemicals are well-represented in the subpallium (**Supplementary Figures 7a–d**). This subpallial appetitive preference mirrors the reported stronger response to mating water compared with skin extract in the juvenile zebrafish subpallium ^30^. In most downstream regions, positive valence was represented by larger numbers of neurons (**Figures 4e, f, Supplementary Figures 7a–d**). In parallel to its preference for spontaneous S-T events in motor control, the pallium also exhibited a more pronounced representation of negative valence compared with other brain regions (**Figures 4e, f**). Of note, although neurons involved in spontaneous S-T event and negative valence representation were outnumbered, they exhibited significantly higher activation levels compared with the counterparts encoding independent tail flips and positive valence, respectively (**Supplementary Figures 8a, b**).

Next, we analyzed the neuronal sensorimotor preferences to understand how individual neurons encode both motor actions and chemical valence (**Figures 4g–k**). For each neuron, we calculated a joint preference measure as the *L*^2^ norm of a two-dimensional vector consisting of the two preference measures (i.e., [motor preference, valence preference], used in **Figures 4c, e**) (see **Methods** for details). We used the measure to quantify a neuron’s preferential responsiveness to either S-T events combined with aversive stimuli, or independent tail flips combined with appetitive stimuli. This analysis further illustrated that the pallium exhibited the strongest overall preferential co-encoding of spontaneous S-T events and negative valence (**Figures 4h, i, Supplementary Figures 8c, d**). In contrast, more caudal brain regions better co-represent spontaneous independent tail flips and positive valence (**Figures 4j, k**).

### Putative pallial involvement in amplifying coordinated eye-body turns

We further examined the temporal dynamics of pallial activity in relation to chemical cue-associated S-T events. Specifically during aversive chemical presentation, pallial motor-encoding neurons exhibited early activation preceding tail flip onset in S-T events, but not during appetitive chemical presentation (**Figure 5a, *upper*, Supplementary Figure 9**). This was also noted in rhombomeres 4 and 6 (**Supplementary Figure 9**), but not in the other brain regions (**Figure 5a, *upper*, Supplementary Figure 9**).

**Figure 5.**
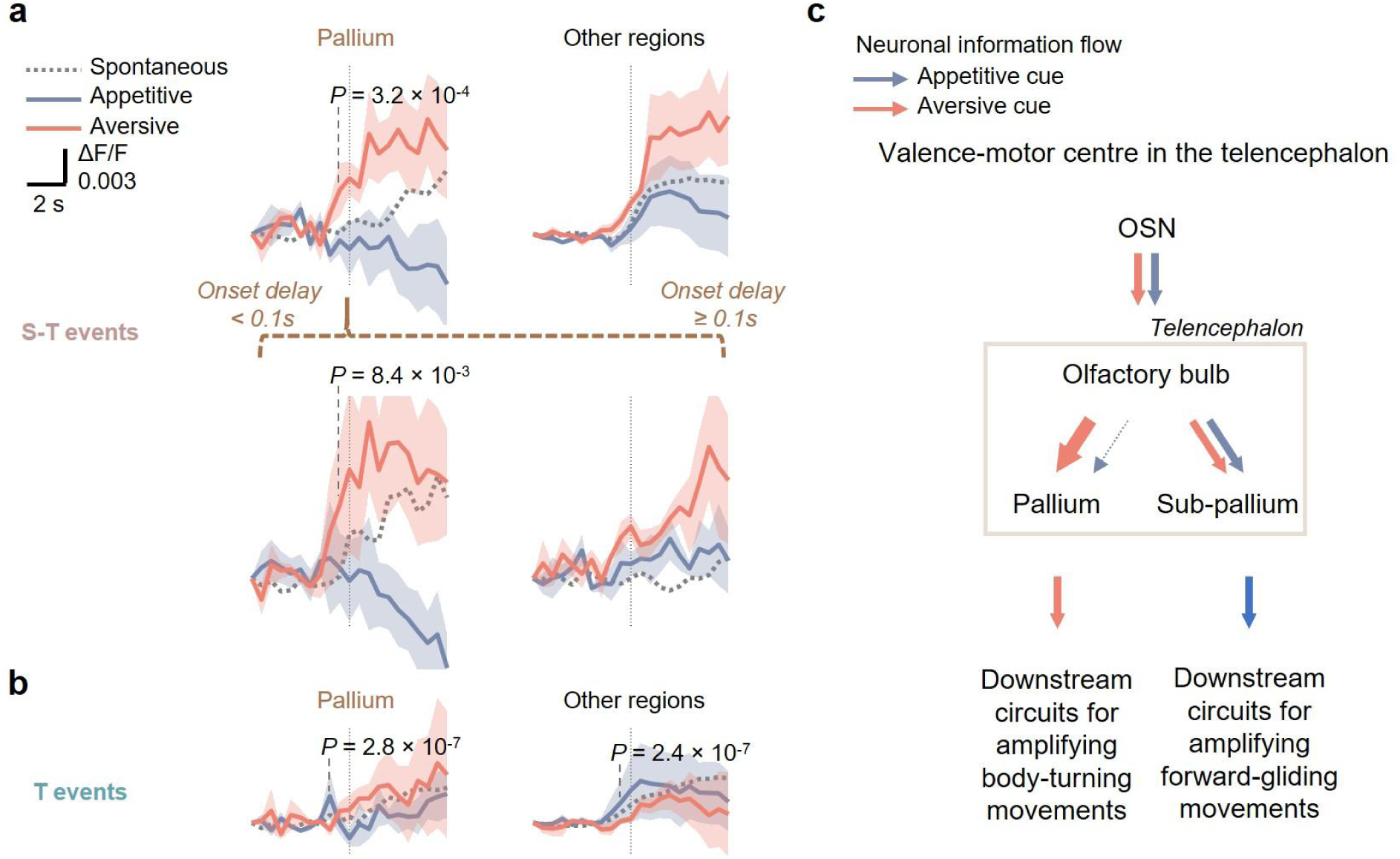
Pallial motor neuron activity onset precedes other brain regions during chemical cue-associated coupled saccade-tail flip (S-T) but not independent tail flip (T) events. **(a)** Upper panel: Mean ± S.E.M. (across larvae, using per-larva averages) responses of motor-encoding ROIs in the pallium (left) and non-pallial brain regions (right) to chemical cue-associated S-T events. Lower panel: Mean ± S.E.M. (across larvae, using per-larva averages) responses of motor-encoding ROIs in the pallium to chemosensory cue-associated S-T events with saccade-tail flip onset delay (tail flip onset-saccade onset) < 0.1 seconds (left) or ≥ 0.1 seconds (right), as defined in Figure 2a. **(b)** Mean ± S.E.M. (across larvae, using per-larva averages) responses of motor-encoding ROIs in the pallium (left) and other brain regions (right) to chemical cue-associated independent tail flips. In **(a)** & **(b)**, dotted traces show the corresponding neuronal responses of spontaneous events. Dotted vertical lines indicate the time points of tail flip onsets (in a corresponding neuronal imaging frame). Neural activity data are from *n* = 5 larvae and correspond to a subset of the behavioral dataset shown in Figures 2 & **3** (*n* = 8). *P*-values: Right-sided *Z*-tests comparing pre-chemical cue-associated event neuronal activity at each of the two time points preceding the motor event to zero. Significance was set at 2.5 standard deviations (S.D.), with S.D. estimated from spontaneous event baseline activity variability. Adjustment for multiple comparisons across time points and brain regions was made using the Benjamini-Hochberg procedure (see **Methods**). **(c)** Schematic illustrations showing the role of pallial activity in linking negative chemical valence with enhanced body-turning movements. Red and blue arrows indicate the proposed schematic relationships associated with processing of negative and positive chemical cue valence, respectively. Abbreviation: OSN, olfactory sensory neurons.

In the telencephalon, pallial motor activations were more prominent for aversive chemical-associated S-T events with a higher temporal proximity of saccade and tail flip onsets (**Figure 5a, *lower*, Supplementary Figure 10a**), and of aversive chemical stimulus and tail flip onsets (**Supplementary Figure 10b**). The first neural activity pattern aligns with the behavioral observation that change in tail flip asymmetry was inversely related to saccade-tail flip onset delay (**Figure 3g, Supplementary Figure 6b**). Interestingly, these were not observed for rhombomeres 4 or 6, where the early motor activations appeared independent of these event proximities (**Supplementary Figures 10a, b**). These observations support an association between pallial motor-encoding activity, the increased tail flip asymmetry and strengthened saccade-tail flip coupling observed during aversive cues. In contrast, the activation patterns in rhombomeres 4 and 6 suggest a premotor role for subsets of neurons in these hindbrain regions, possibly contributing to S-T events independently of their temporal proximity to the initiation of aversive chemical stimulus.

In contrast to S-T events, independent tail flips exhibited distinct associated neuronal activity patterns. These were characterized by a lack of early pallial activation relative to tail flip onset during aversive cues presentation (**Figure 5b, Supplementary Figure 11**). Instead, independent tail flips were associated with early neuronal activations in the preoptic area, pretectum, tectum, rhombomere 7, and caudal hindbrain, particularly during appetitive stimuli (**Supplementary Figure 11**). This differential activation pattern further underscores the role of pallium in representing S-T events, especially in response to aversive stimuli.

Collectively, these findings align pallial activity with the aversive chemical-specific enhancement of saccade-tail flip coordination observed behaviorally (**Figure 3g**). They also highlight the larval zebrafish pallium as a prominent sensorimotor locus where valence-specific chemosensory and motor-related signals converge (**Figure 5c**). While these activity patterns strongly track valence-dependent changes in coordination, we acknowledge that our data are correlative and do not establish that pallial circuits are causal in this modulation, which requires additional perturbation experiments to prove.

## Discussion

Chemosensation is one of the most ancient and conserved sensory mechanisms, playing a critical role in guiding locomotion and body orientation across the animal kingdom^26^. While its influence on body movements is well-documented, our study provides the first direct evidence that chemosensory inputs modulate eye-body coordination, a dynamic interplay previously thought to rely primarily on visual and mechanosensory cues. Using larval zebrafish as a model system, we uncovered a sensory-motor integration mechanism wherein chemosensory valence shapes not only locomotor behaviors but also the temporal and directional coupling of saccades and body turns. These findings reveal a previously unknown synergy between oculomotor and chemosensory systems, expanding our understanding of how multisensory integration drives adaptive motor control.

Larval zebrafish swim and make body turns in discrete bouts, a behavior that reflects a foundational motor pattern common in many animal species during development ^67^. We first show that saccades and tail flips are tightly coupled in time and direction for peak body turn. This coordination likely optimizes visuomotor behaviors such as tracking and orienting, which are essential for survival. Chemosensory valence further modulates this coordination: aversive cues enhance saccade-tail coupling, associating with sharper turns and more frequent saccade-coupled movements, which may have a role in updating the gaze along with larger reorientation and preparing the visual system for updated input from the new heading; appetitive cues promote forward gliding with minimal impact on saccade-tail coupling. In this view, chemosensory signals operate on a slightly slower timescale, modulating the likelihood and amplitude of coordinated orientation and gaze shifts, while visual cues provide fast, precise guidance for individual movements, allowing integration of modalities with different temporal dynamics. These distinct motor programs may facilitate context-specific responses, such as more efficient predator evasion, or foraging in favorable environments. This modulation of visuomotor coordination by chemosensory input aligns with conserved principles of motor control described in other vertebrates, such as in *Xenopus laevis*, where eye and body movements are synchronized via feedforward efferent copies of spinal central pattern generator activity. Notably, work in Drosophila has shown that visual and olfactory cues jointly shape steering and odor-tracking ^68,69^, suggesting that reorientation and active visual sampling are common multisensory strategies in small animals. In general, such multi-modal information integration should help to ensure gaze stability and locomotor efficiency.

Because the observed behavioral effects were largely confined to the 10-s stimulus window and returned to baseline thereafter, they are consistent with a rapid, cue-locked modulation of eye–body behavior rather than a sustained change in internal state. However, this temporal profile does not exclude a transient arousal-related contribution. In particular, aversive cues increased saccade frequency, which could increase the opportunity for threshold-defined saccade–tail-flip co-occurrence. However, circular saccade-time shuffling did not reproduce the temporal and directional organization of aversive-context S-T events, indicating that altered event rates alone cannot explain their coordinated structure. Experiments that independently manipulate arousal or saccade and tail-flip frequencies will be required to resolve their contributions to S-T event prevalence and coordination.

At the neuronal circuit level, our whole-brain imaging points to the telencephalic pallium as a forebrain node where negative chemosensory context and S-T-related motor signals converge, providing a candidate neuronal substrate for valence-dependent modulation of eye–body coordination. Negative chemical valence was strongly represented in olfactory circuits and downstream relay regions, suggesting that aversive cues engage specific sensorimotor pathways associated with enhanced saccade-coupled turns. In line with this, a recent study has shown that transcriptionally defined domains and a tyrosine hydroxylase-expressing interneuron population in the larval olfactory bulb are differentially recruited by attractive versus aversive odors and are causally required for attraction but not aversion, indicating that valence-specific transformations emerge already at the level of the olfactory bulb ^70^. Complementary imaging work in larval zebrafish has mapped chemosensory valence onto distributed neuronal populations across forebrain regions, including pallium, under matched appetitive and aversive cues ^71^. The pallium, an evolutionarily conserved dorsal telencephalic structure in vertebrates ^64^, has been implicated in diverse functions, including sleep regulation ^72^, place encoding ^73^ and spatial representations ^74^, and was recently shown to participate in hierarchically organized thalamocortical-like sensory processing ^75^, as well as visually guided social approach behavior ^76^. Consistent with a broader role in aversive processing, another recent work has identified negative-valence-selective neurons in the larval zebrafish pallium that respond preferentially to strongly aversive stimuli ^77^. Our findings extend these observations by showing that pallial activity reflects negative chemosensory context but also tracks the likelihood of S-T events at an early developmental stage. However, because we did not perturb pallial activity, we interpret their role as correlative and predictive for S-T events, whereas a general causal motor command role will need further experimental proof.

The evolutionary implications of these findings are significant, indicating that the ability to integrate chemosensory input with motor programs likely emerged over 450 million years ago ^78^. By revealing how chemosensory valence modulates eye-body coordination, our study sheds light on the conserved nature of sensory-motor integration. This mechanism may be present across other vertebrates, though species-specific adaptations—such as the absence of necks in teleost fish ^44,45^—likely shape and determine species-specific patterns of coordination. In mammals or birds, for instance, head and neck movements may play a more prominent role in sensorimotor integration.

This work opens avenues for further studies on sensory input-mediated eye-body control. First, the neuronal circuits underlying chemosensory modulation of eye-body coordination should be explored in greater detail. High-resolution imaging of specific brain regions, such as the pallial and subpallial structures, could reveal how chemosensory valence is encoded and linked to motor commands. Second, extending these studies to other species, including vertebrates with flexible necks, could provide insights into how evolutionary pressures have shaped sensorimotor integration. Third, investigating additional sensory modalities (e.g., auditory ^79^) and their interactions with chemosensation may uncover more general principles of multisensory integration.

In conclusion, our findings reveal a new mechanism by which chemosensory cues modulate eye-body coordination, providing insights into the neural basis of adaptive locomotion. By integrating behavioral and neuronal findings within an evolutionary framework, this work advances our understanding of sensory-motor integration and lays the foundation for future studies on multisensory control of motor behaviors.

## Key resources table

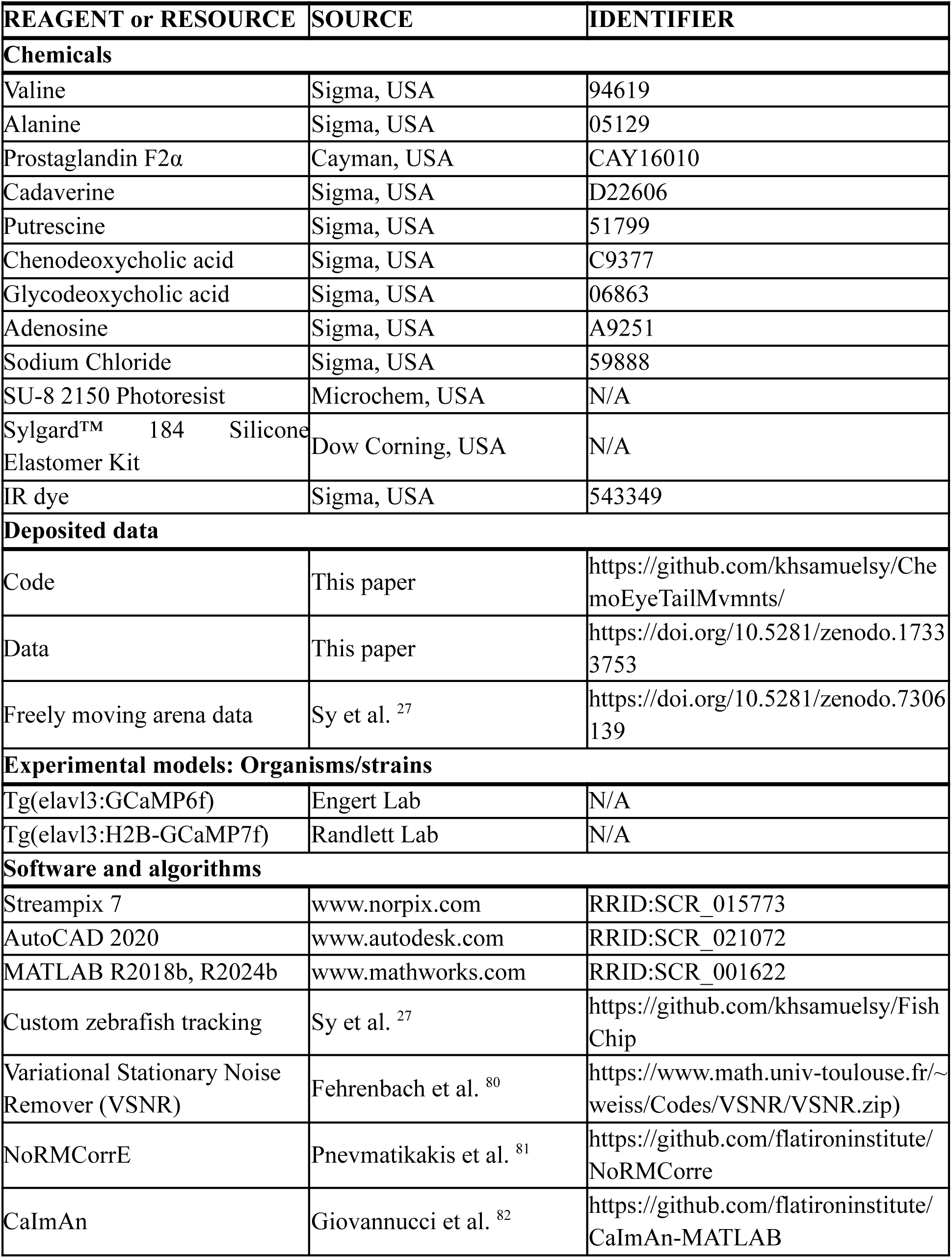
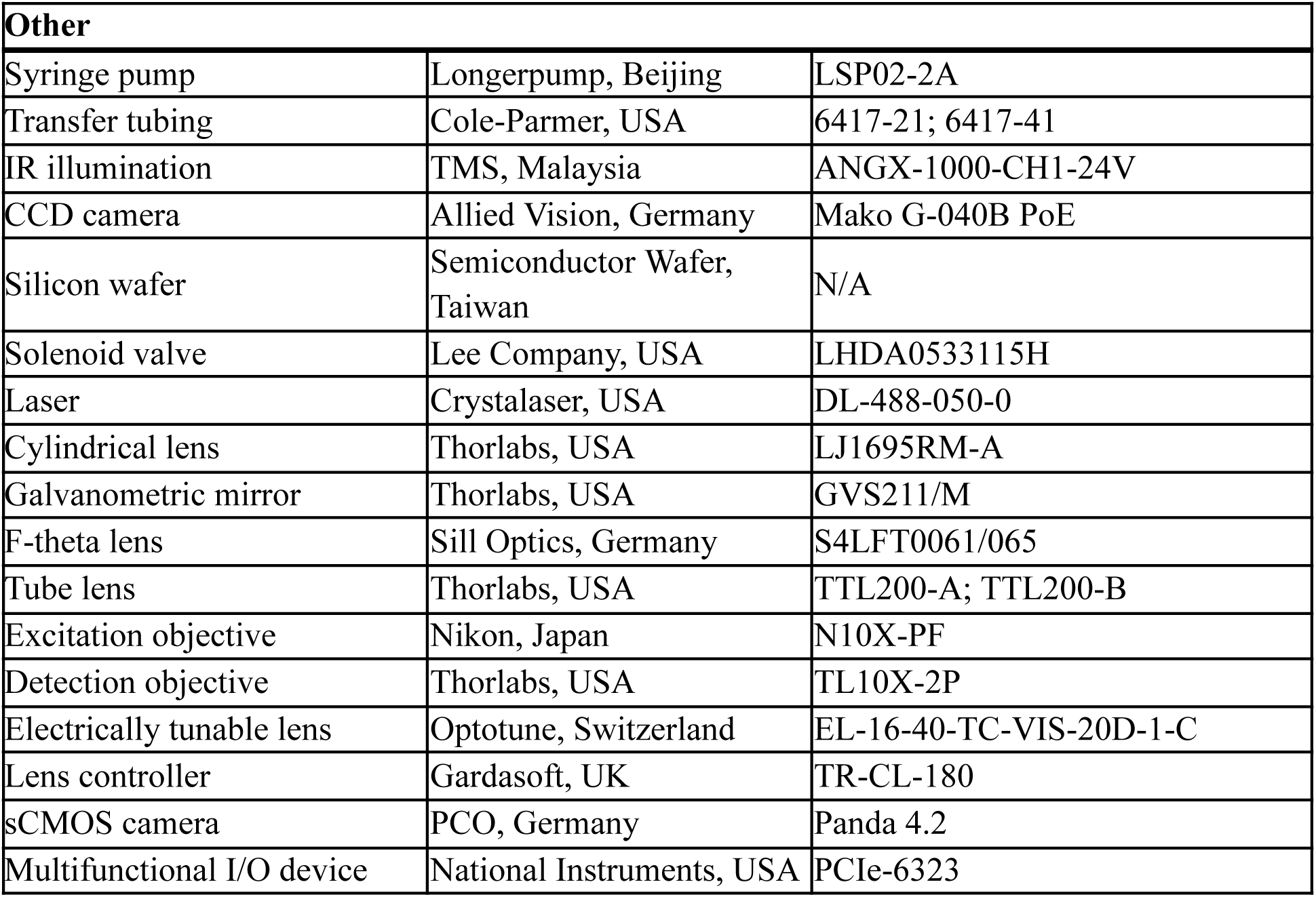

## Methods details

### Animals

All experimental procedures received prior approval from the Animal Research Ethical Committee of the Chinese University of Hong Kong (CUHK) and complied with the Guide for the Care and Use of Laboratory Animals. For this study, *nacre*-mutant larvae, aged 5–8 days post-fertilization (d.p.f.), were raised and maintained under a 14-hour light/10-hour dark cycle at a constant temperature of 28°C. Larvae used for experiments were not fed, as per common practice and this does not compromise their health at this developmental stage^83^. The larvae used in the valence determination assay (*n* = 313) were bred from adult zebrafish pairs with at least one heterozygous Tg(elavl3:GCaMP6f) parent and were not screened for GCaMP6f expression. The larvae used in both the behavioral and neuronal imaging assays (*n* = 5) were bred from Tg(elavl3:H2B-GCaMP7f) adults and were screened for the presence of GCaMP7f expression. Larvae used exclusively for behavioral imaging (*n* = 3) were derived from the same line but were not screened for expression. The sex of larvae was not determined at this stage of development.

### Fluidics-based swimming arena for chemical valence determination

The device was designed using AutoCAD 2020 (Autodesk, USA) and comprised five 1.5 mm-thick layers of laser-engraved poly(methyl methacrylate) (PMMA). The layers were meticulously aligned vertically and fused by applying a chloroform solution along the contact edges, which yielded a durable, sealed structure. This process formed a closed square arena with dimensions of 40 mm in length and 1.5 mm in height (**Figure 1a**). An additional inlet was used for loading larvae and sealed prior to assays. The inlets and outlets are 200 µm wide, prohibiting larvae from exiting the arena through these channels. Fluidic connections were established by securing syringe needles (Terumo, USA) to the topmost PMMA layer with an epoxy adhesive paste (Devcon, USA). These needles were linked to the inlet and outlet channels, which in turn connected to syringes containing the fluid stimuli and to a waste collection bottle, respectively. The connections were facilitated using poly(tetrafluoroethylene) (PTFE) tubing (Cole-Parmer, USA). After construction, the device was left to air-dry for 24 hours, allowing for the complete evaporation of residual chloroform and epoxy adhesive paste. To ensure the device was contaminant-free, it was washed thoroughly with water twice, thus removing any potential contaminants prior to its use in behavioral assays.

Defined chemical zones within the arena were created using four fluid streams, each channeled through inlet channels located at the corners of the arena and propelled at a constant flow rate of 100 µL/min by syringe pumps (LSP02-2A, Longerpump, Beijing). The background solvent of these fluid streams was either water or a 0.5% DMSO solution, depending on the tested chemical species. In total, 9 chemicals were tested, including valine (Val; Sigma, USA), alanine (Ala; Sigma, USA), prostaglandin F2α (PGF2α; Cayman, USA), cadaverine (Cad; Sigma, USA), putrescine (Put; Sigma, USA), chenodeoxycholic acid (CDCA; Sigma, USA), glycodeoxycholic acid (GDCA; Sigma, USA), adenosine (Ade; Sigma, USA), and sodium chloride (NaCl; Sigma, USA). Chemicals with limited water solubility, including PGF2α, CDCA, and GDCA, were dissolved in 0.5% DMSO solution to ensure complete dissolution. This concentration of DMSO has been determined to be safe for the larvae and does not affect their baseline behaviors ^84^. Other chemicals were dissolved in water.

The top right control quadrant (quadrant I) contains no chemicals, while the remaining quadrants contain chemical species at varied concentrations (**Figures 1a, b**). For the three chemical species with limited solubility, both water and 0.5% DMSO were used as controls in separate assays (**Supplementary Figures 1d, e**). The outlet of the arena was connected to an open waste collection bottle to allow for the efficient disposal of the outflow fluids.

The Reynolds number (*Re*) of fluid flow in the arena is given by *Re* = *⍴vD*_H_*/μ*, where *⍴* is the fluid density, *v* is the fluid flow velocity, *D*_H_ is the hydraulic diameter of the device, and *μ* is the fluid dynamic viscosity. Given the average cross-sectional dimensions at each quadrant of the arena (i.e., 1.5 mm × 10√2 mm), the corresponding average flow rate was 100 μL min^-1^ or 1.67 mm^3^ s^-1^. Hence the average flow velocity = 1.67 mm^3^ s^-1^/(1.5 mm × 10√2 mm) = 7.86 × 10^-5^ m s^-1^. *D*_H_ for the rectangular channel is given by 2*ab*/(*a*+*b*) = 2.71 × 10^-3^ m, where *a* and *b* are the average dimensions of the rectangular cross-sections (i.e., 1.5 mm and 10√2 mm). Substituting the *D*_H_ found, and the density (996 kg m^-3^) and the dynamic viscosity (8.32 × 10^-4^ Pa.s) of water at 28 °C, the Reynolds number was found to be *Re* = 0.255 << 2000. As *Re* << 2000, the streams were laminar with negligible mixing ^85^ and static fluid zone borders were formed. The Péclet number (*Pe*) of fluid flow in the arena is given by *Pe* = *vD*_H_/*D*, where *D* is the diffusion coefficient of the molecule. Since all the investigated small molecules have diffusion coefficients of the 10^-9^ m^2^s^-1^ range order, *Pe* ≈ 2.13 × 10^2^ >> 1, the molecules thus have minimal diffusivity across the streams. The *Re* and *Pe* values suggest that the chemicals delivered using the system would be minimally crossing via advection and diffusion, respectively, and therefore stay highly localized within individual fluid streams. As the geometry and flow profile of the four quadrants are identical, four identical static chemical zones are established. The chemical zones were visualized by infusing an IR dye (IR 806, Sigma, USA) into the quadrant II and IV of the device (**Figure 1a, *lower***).

### Chemical valence determination assay

During a chemosensory valence determination assay, three to six zebrafish larvae aged 5–8 days post fertilization (d.p.f.) were placed in a swimming arena and allowed to acclimate for 15 minutes before the introduction of chemical stimuli. To minimize potential visual cues across different assays, the arena was horizontally rotated to a single angle of 0°, 90°, 180°, or 270° in a pseudo-randomized manner for each assay. To further reduce visual cues, the recording was conducted under 850-nm infrared illumination provided by a backlight LED panel (BHS4-00-100-IR850-24V, TMS, Malaysia) controlled by an ANGX-1000-CH1-24V driver (TMS, Malaysia), in the absence of visible light. Time-lapse video was recorded by a CCD camera (Mako G-040B PoE, Allied Vision, Germany) at 20 frames per second, with image acquisition managed by Streampix 7 (Norpix, Canada). The recording lasted for 30 minutes, with four fluid streams flowing at the same speed. The fluid temperature was maintained at ∼28°C throughout the assay. This assay enables precise behavioral monitoring of the larvae, ensuring that the specific chemical exposure was exactly known, thereby minimizing potential biases in hedonic measurements.

### Microfluidic device for multichannel precise chemical stimulation, saccade, tail flip and brain-wide cellular-resolution imaging

Capturing the behaviors and neural activity elicited by chemosensory cues presents a significant challenge in the field. In our previous development of Fish-on-Chips ^27^, we enhanced the precision of a single chemical cue delivery to larval zebrafish, enabling stable recordings of neuronal responses to chemosensory inputs. In this work, we further developed the platform with the objective of systematically exploring the impact of multiple ecologically pertinent, valence-inducing chemicals on the natural behaviors of larval zebrafish and uncovering the associated neural circuitry.

Compared with our previous device, we further optimized the channel design in order to achieve multiple chemical delivery, compatibility with dual-scanning light sheet imaging and maximize the behavioral readout to include both saccade and tail flip. The PDMS-based microfluidic device was designed in AutoCAD 2020 (Autodesk, USA). It was then fabricated by customized photolithography techniques like the previous ones to incorporate flat transparent sidewalls at the front and the right side with respect to the head chamber to accommodate for the entry of two scanning excitation light sheets, and a thin glass ceiling for the emitted fluorescence detection path. Briefly, a 2D design was printed on a soda lime mask (Supermask, Shenzhen). Negative photoresist SU-8 2150 (Microchem, USA) was spin-coated on a 4-inch silicon wafer (Semiconductor Wafer, Taiwan) to 500-μm thick. The device pattern was then transferred to the photoresist with UV exposure (OAI, USA), followed by post-exposure bake and etching to produce a master mold. Mixed PDMS (Dow Corning, USA) and curing agent (at 10:1 w/w) were poured into the salinized mold to a thickness of 5 mm, with perpendicular glass slides held upright on the mold to produce the front and the right side flat vertical surfaces upon curing. After solidification at either 65°C for 240 minutes or 100 °C for 60 minutes, it was detached from the mold, bonded and sealed onto a microscopic cover slide (as thin glass ceiling) by plasma treatment. All devices were rinsed with water twice before use in experiments.

The fluidic stream control solution was similar to our previous design. In brief, bilateral three-layered, sandwiched fluid streams are directed to the larval zebrafish from the sides and converge at the middle, and then leave the device. The middle stream was switched to different solutions to allow selections of chemicals in different trials. Each fluid stream was carried by PTFE tubes, controlled by a solenoid valve (LHDA 0533115H, Lee Company, USA), and driven by a syringe pump. A side channel connecting the chemical delivery microchannels and the tail chamber was incorporated to buffer pressure changes in the fluidic environment, and minimize mechanical disturbances to the larval subject. The flow rate was uniformly adjusted at 80 µL/min, which yields the same order magnitude of *Re* and *Pe* numbers, and thus allows for highly controlled fluid delivery. The details of the valve control and chemical delivery are described in the following sections.

Through iterative experimentation, we determined the necessity for bilaterally independent channels to achieve symmetrical stimulus rise and fall times. One challenge we noted was the natural formation of micro-sized air bubbles during fluid coalescence, which can interfere with the accuracy of fluid delivery. The likelihood of bubble formation scales with the number of fluid inlets. We optimized the design to include a total of 7 channels on each side, using the first and last channels for water streams (sandwiching the chemical streams), and 5 channels dedicated to different chemical streams (**Figure 1d**).

These significant enhancements, together with other minor ones, enabled us to systematically deliver 4 different water-soluble chemicals plus 1 blank water control for 4–6 repeated trials each. Appetitive and aversive chemosensory stimuli were delivered as brief trials and presented in a balanced, pseudo-random order, so that neither cue type was presented continuously for extended periods and appetitive trials were interleaved with aversive trials, with each larva experiencing all stimulus types (blank, appetitive, and aversive conditions) in this within-subject design. We ensured a 10-second exposure duration for each stimulus trial to allow sufficient time for the larval subjects to exhibit saccadic and tail flipping responses, which differed greatly in event frequency (**Figure 1e**). Additionally, we also ensured a 10-second imaging time after each stimulus was removed. This setup was optimal for multi-trial simultaneous behavioral-neuroimaging experiments, allowing for approximately one hour total assay time per larval subject. We anticipate incorporating additional channels and an even more streamlined experiment paradigm with further developments.

### Dual-scanning light sheet fluorescence microscope for whole-brain calcium imaging

We custom-built a light sheet fluorescence microscope with two light sheet excitation paths from the front and right side of the larval subject for optimized cellular-resolution whole-brain imaging. For each of the excitation paths, a 486 nm-centered blue gaussian laser (DL-488-050-0, Crystalaser, USA) was used as the excitation light source. The laser beam was resized to 0.6 mm in diameter (1/*e*^2^) by a pair of telescopic lenses (LB1757-A, LB1596-A, Thorlabs, USA), which then passed through a scanning system, followed by a cylindrical lens (LJ1695RM-A, Thorlabs, USA) that focused the horizontal dimension of the parallel beam onto the back focal plane of an air excitation objective (Nikon Plan Fluorite, ×10, N.A. 0.3, 16 mm WD). This expanded the laser horizontally to form a light sheet. The scanning system consisting of a galvanometric mirror (GVS211/M, Thorlabs, USA), a F-theta lens (S4LFT0061/065, Sill Optics, Germany) and a tube lens (TTL200-A and TTL200-B, Thorlabs, USA) was used to scan the light sheet vertically and linearly over a range of 276 µm. The F-theta and tube lenses also expanded the beam 3.31 times, resulting in a beam diameter of 2 mm (1/9^th^ of the objective back aperture) and effective excitation N.A. of 0.0332. A photomask with a 250 µm aperture and a photomask with an open-ended aperture (**Supplementary Figure 2c**) were placed and aligned at the front and at the right side of the device respectively to avoid directly shining the eyes of the larval subject with either light sheet at either side.

Along the detection path, a bandpass green filter (525 ± 25 nm) was placed after an air detection objective (TL10X-2P, Thorlabs, USA; ×10, N.A. 0.5, 7.77 mm WD) to block the blue excitation light. An electrically tunable lens (EL-16-40-TC-VIS-20D-1-C, Optotune, Switzerland) was linearly driven by a lens controller (TR-CL-180, Gardasoft, UK) and synchronized with the light sheet scanner to achieve rapid focusing of different image planes onto the sensor of a sCMOS camera (Panda 4.2, PCO, Germany). During an experiment, to correct for axial drift, a reference plane was calibrated with respect to the initial measurement at the beginning of each trial. All control units were synchronized using a multifunctional I/O device (PCIe-6323, National Instruments, USA) and custom-written codes in MATLAB (R2018b, MathWorks, USA).

### Simultaneous whole-brain calcium imaging and behavioral recording

To obtain the most naturalistic data, we only included larvae that were nearly perfectly leveled in the device, as evidenced by both eyes being at the same vertical level, for further experimentation and analysis. There were no further exclusions of animals. In total, 8 larvae (5 at 6 days post fertilization (d.p.f.) and 3 at 5 d.p.f.) drawn from 6 breeding batches, were used. Of these, 5 larvae underwent simultaneous behavioral and whole-brain neuronal calcium imaging experiments in an upright orientation as permitted by the improved imaging set-up, including 4 at 6 d.p.f. and 1 at 5 d.p.f., while the remaining 3 underwent behavioral experiments only.

In an imaging experiment, a larva was loaded (via the larva inlet) and fitted into the trapping chamber of the PDMS-based microfluidic device using a syringe pump and monitored under a surgical microscope. Then, the larva was allowed to acclimate for 15 minutes. Introduction of bubbles to the microfluidic system was carefully minimized during system setup to avoid unwanted fluid flow pattern changes or mechanical instability of the larval subject. Prior to functional imaging, a detailed anatomical stack of the larval zebrafish brain spanning 138 imaging planes at 2-µm intervals was taken. In this intact upright imaging configuration, the majority of the brain except the topmost (e.g., habenula) and bottommost regions (e.g., caudal hypothalamus) which are directly under the cover glass slide and furthest from the detection objective lens, was consistently imaged across larvae.

During functional imaging, whole-body saccadic and tail flipping behaviors (single-plane, at 200 Hz) and volumetric whole-brain calcium fluorescence signals (28 planes with 7-µm intervals, at 2 Hz) were simultaneously imaged under chemosensory cue (or blank control) delivery. Behavioral images were acquired under IR illumination by a CCD camera (Mako G-040B PoE, Allied Vision, Germany) and the Streampix 7 (Norpix, Canada) image acquisition software, while calcium images were acquired with the abovementioned custom dual-light sheet scanning microscope. Each larva underwent 23–30 experimental trials with a total of five types of stimuli. These stimuli include amino acids (valine and alanine), diamines (cadaverine and putrescine), all at one millimolar concentration, and a blank control. Each stimulus was delivered bilaterally to both nostrils for 4–6 repetitions. Each trial lasted for 50 seconds with 30-second baseline, 10-second stimulus presentation, and 10-second post-stimulus recording periods, followed by ∼90-second inter-trial resting intervals. Axial drift was corrected between trials for larvae undergoing simultaneous behavioral-neuronal imaging.

During each 30-second baseline recording period, in the three-layer sandwiched fluid streams, the middle stimulus stream carried a chemical cue (or water for blank control trials). The first (water) and the second (stimulus) stream in front of the nostrils were then switched off sequentially by the respective controlling valves at the two sides, with 10-second gap to allow the second (stimulus) and the third stream (water) to contact the nostrils. This resulted in the initiation and termination of chemical stimulation. Recording continued for 10 seconds after stimulus cessation.

### Valence determination analysis (swimming arena)

Individual frames of the navigation behavioral tracking videos were first registered for translation, background-subtracted, and contrast-adjusted. The *xy*-coordinates of the heads of larvae in each frame were extracted using our previously established semi-automated template matching-based tracking program ^27^ custom-written in MATLAB (R2018b, MathWorks, USA). All tracking results were manually verified on a frame-by-frame basis and corrected when necessary. The coordinates were then used to calculate the proportions of time spent in each of the same-area quadrants over the 30-minute recording time. Additionally, since there is minimal mixing across streams under laminar flows, interactions between quadrants containing the same chemical at different concentrations were negligible. The population standard deviation of the time spent difference in chemical quadrants (compared with the control quadrant) of the setup was estimated from the sample standard deviation of the time spent difference in the control group with Bessel’s correction.

### Behavioral image processing and analysis (microfluidic device)

Eye and tail movements were extracted from behavioral images by a custom MATLAB program (R2018b, MathWorks). The first ten seconds of the behavioral recordings of each trial were excluded from analysis to avoid confounding behaviors evoked by the onset of light sheet illumination. In addition, a total of six trials, three trials each from two subjects, were found to have bubbles present, which disrupted the laminar flows necessary for precise chemical delivery. These trials were also discarded. For eye movements, the video was down-sampled by a factor of 25 times (resultant frame rate: 8 fps) to achieve a higher signal-to-noise ratio. The turn direction and duration of the saccades were manually identified using rotational visual flows in each region-of-interest of the two eyes. Only 5 events (out of 843 events recorded from 8 larvae) of bilateral eye movements from 3 larvae were recorded with incoherent eye movement directions, and these were excluded from analysis. Pre-saccade interval was defined as the time between the onset of a saccade and that of its preceding one, if both were recorded.

For tail movements, episodes of tail flips were identified based on changes in tail tip-waist angle using several criteria. An episode was isolated when tip-waist angle changes occurred in at least two consecutive frames (frame rate: 200 fps), and were temporally situated before and after at least three frames of stationary angle changes. The start and end frames were determined as the last frame of the last idle period and the first frame of the next idle period, respectively. Positive and negative peaks in the undulating cycles of angle changes were identified using the built-in *findpeaks* function in MATLAB (R2018b, MathWorks, USA). The undulation cycle was determined by averaging the number of positive and negative peaks.

Pre-tail flip interval was defined as the time between the onset of a tail flip and that of its preceding one, if both were recorded. Tail flip duration was calculated as the time difference between the end and start frame of an episode. The average undulation was calculated by dividing the duration by the undulation cycle. Magnitude and tail flip asymmetry were calculated using the sum and difference of the average net positive and average net negative peaks, respectively, with the zero reference being the average of the start and end angles (**Supplementary Figure 3d**). A small non-zero average tail flip asymmetry to the right side was consistently observed in the spontaneous tail flips of the five subjects undergoing simultaneous behavioral-neuronal imaging (2.6° ± 1.2° (mean ± S.D.)), which was due to the asymmetrical illumination from the right side. This bias was substantially smaller in the three subjects which underwent behavioral imaging only and were not subject to ambient excitation light (0.6° ± 1.0° (mean ± S.D.)). Tail flip asymmetry calculation was adjusted by subtracting each subject’s average tail flip asymmetry before further analysis.

Tail flips were classified based on tail flip asymmetry. A tail flip with an asymmetry of ≥ 1.5° (one-half of the median asymmetry of all spontaneous tail flips) was considered a turn event. If the asymmetry was < 1.5°, it was classified as a non-turn event. In the laterality analysis of tail flips (**Figures 2f, h, Supplementary Figures 3c, 5e, f**), only turn events were included for comparisons. Non-turn events, indicating forward movement, were excluded from this analysis.

A tail flip was defined as saccade-coupled (S-T events) if the onset occurred within 0.5 seconds of a saccade’s onset (i.e., |T-ON - S-ON| ≤ 0.5 seconds). For each saccade, only the tail flip with the closest onset was considered saccade-coupled. S-T events with saccade onset certainly later than the tail flip onset, defined as those with T-ON - S-ON < -0.0625 seconds with the temporal error of saccade time determination considered, represented a small subset (28 out of 249) of all spontaneous S-T events (**Figure 2b**). This pattern was largely conserved for chemical-associated S-T events (appetitive: 8 out of 46, aversive: 3 out of 57, **Supplementary Figures 5a, c).** These were not included from the PCA and regression analyses of S-T coordination (**Figures 3g, h, Supplementary Figure 6**).

Events recorded during blank trials were classified as spontaneous. During chemical stimulation trials, events were considered spontaneous if the onset was recorded before the stimulus onset, and the offset was recorded before or coincided with the stimulus onset. Events were considered to be stimulus-evoked if the onset occurred later than stimulus onset, and the offset occurred before or coincided with the stimulus offset. Onset and offset of S-T events were defined as that of the saccade. Events during the resting intervals between trials were not recorded or analyzed.

Pre-event interval was defined as the time difference between the onset of a saccade, a tail flip, or a bout and that of the respective preceding one (**Figures 1e, 2d, e, Supplementary Figure 4a**). For comparison with flanking events (**Figures 2d, e, g–j, Supplementary Figures 3e, f**), saccades and tail flips were first categorized into either S-T or independent groups and set as the 0^th^ next event. For each 0^th^ next event, the sequence and quantity of other events (n^th^ next event) were shown. An event could be the 1^st^ next event for one sequence and the 2^nd^ next event for another, causing overlaps.

### Comparative analysis based on previously acquired data of freely swimming larvae

A dataset from our previous study was used ^27^. In that study, 11 larvae in 3 assays were allowed to swim in the absence of visible light for 2 hours in a 2D rectangular arena (60 mm × 30 mm × 1.5 mm) completely filled with still water, and imaged under IR illumination. The heading orientation (10° resolution) of every larva in each frame was extracted using a semi-automated template matching-based tracking program custom-written in MATLAB (R2018b, MathWorks, USA). All bouts subsequent to the first bout in the rightmost virtual chemical zone of the arena were extracted for analysis. Pre-bout interval was defined as the time difference between the onset of a swim bout and that of its preceding bout. Turn angle was defined as the orientation change in either direction. A swim bout was defined as a turn bout based on the turn angle being ≥ 70°, and non-turn bout otherwise. Only swim bouts with an orientation change were used to compare the proportion of consecutive ipsilateral turns (**Supplementary Figure 4b**). Because this assay was optimized for multi-animal tracking over a large field of view (60 mm × 30 mm; approximately 0.1 mm per pixel), the larval eyes occupied only a few pixels, and eye rotations could not be quantified with sufficient precision; therefore, eye movements were not analyzed in this dataset.

### Calcium image processing and signal extraction

We adopted an analysis pipeline similar to our previous work ^27^. Detailed larval zebrafish brain anatomical stacks were registered to the Z-Brain Atlas ^86^ using affine transformation followed by non-rigid image registration using custom scripts written in MATLAB (R2018b, MathWorks, USA). Functional imaging planes were matched to the corresponding anatomical stack by maximization of pixel intensity cross-correlation and manually verified. Stripe artifacts in the anatomical stacks and functional imaging frames were removed using the Variational Stationary Noise Remover (VSNR) algorithm ^80^. Functional imaging frames were motion-corrected using discrete Fourier transform and NoRMCorrE algorithms ^81^. Trials with blurred frames due to in-frame drifts were discarded. Region of interest (ROIs) corresponding to individual neurons were then extracted using the CaImAn package ^82^. Anatomical landmarks and the regional identity of each ROI were verified by manual inspection (by S.K.H.S.). Note that this only included neurons that had fired at least once during all imaging sessions and with a sufficiently high signal-to-noise ratio determined by the CaImAn-based analysis pipeline.

Trials with unsatisfactory image registration were excluded from further analysis. For one larva with more jitters in the functional images, the image stacks were initially downsampled to 1 Hz by averaging every two consecutive frames. For trials with plane-specific jitters, a small proportion of frames with jitters were replaced with the immediate preceding frame (or the next frame if the preceding frame also had jitter) for the data from all larvae. Trials with more than two consecutive jittering frames were excluded from further analysis. In total, we retained whole-brain neuronal imaging data from 88 out of 139 trials from the 5 larvae that passed all quality control criteria.

### Identification of motor- and sensory-coding neurons

The first 10.5 or 11 seconds (21 or 11 imaging frames) of the imaging data of each trial were excluded from analysis to avoid the inclusion of transient activity evoked by the onset of light sheet illumination. The minimum fluorescence intensity in the subsequent 4.5 or 4 seconds (9 or 4 imaging frames) of each ROI was used as the baseline fluorescence (F). The ΔF/F signals of the remaining imaging frames were calculated as the calcium response of each ROI.

Motor and sensory regressors were designed to identify motor- and sensory-encoding ROIs. Motor output was defined to be the proportion of frames with active tail flip in each 0.5-s or 1-s imaging time bin, and a motor output regressor was created to represent this (motor regressor 1). A second motor regressor for independent tail flips was created by excluding all tail flips of S-T events (motor regressor 2). A third regressor was designed by randomly removing tail flip events from motor regressor 1 at a probability that corresponded to each larva’s S-T event proportion (motor regressor 3). Utilizing these three regressors allowed us to directly identify both the saccade-coupled and independent tail flip-encoding neurons but avoided saccade-encoding neurons ^6,87^. Chemical stimulus regressors were created with a step function during the stimulus window of each chemical. All regressors are normalized to the [0, 1] range. We calculated the mutual information (*I*) between the calcium responses of each ROI and the regressors by a method based on kernel density estimation of the probability density functions of variables ^88^. For motor mutual information, only all the imaging frames in the entire blank trials and before stimulus onsets in stimulus trials were used for calculation. For sensory mutual information, all imaging frames were used. With the three motor (**Table 1**) and five chemical stimulus regressors, each ROI had eight mutual information values, namely *I*_tail_flip_all_ (based on motor regressor 1), *I*_tail_flip_independent_ (based on motor regressor 2), *I*_tail_flip_randomRemove_ (based on motor regressor 3), as well as *I*_valine_, *I*_alanine_, *I*_cadaverine_, *I*_putrescine_, and *I*_blank_ (for which the suffix denotes the chemical or blank control). These values were used to determine significant motor and/or sensory information encoding, and calculate the motor and valence preferences (see the next sections).

**Table 1.**
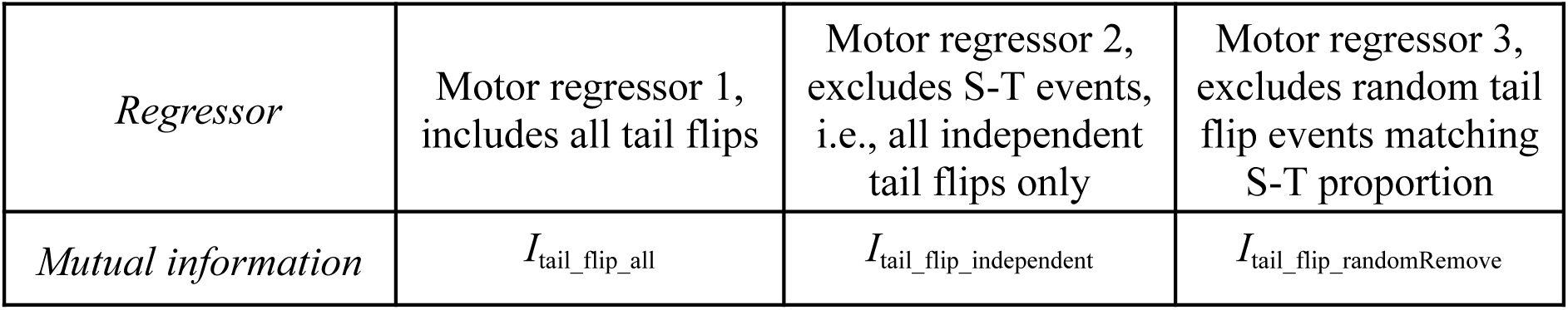
List of motor regressors and mutual information measures.

To estimate the distributions of the mutual information values expected under randomness, we shuffled the ΔF/F signals of each ROI (i.e., disrupting their temporal structures) and calculated another set of shuffled values. We defined the motor- and sensory-encoding neurons to be those that most significantly encode behavior variables or chemical stimuli. Motor-encoding neurons were defined as ROIs with either *I*_tail_flip_all_ or *I*_tail_flip_independent_ > 1.3 times 99^th^ percentile of the shuffled *I*_tail_flip_all_ or *I*_tail_flip_independent_, respectively. Sensory-encoding neurons were defined as those with at least one *I*_chemical_ > 1.3 times the 99^th^ percentile of the shuffled *I*_chemical_ (where the suffix refers to one of the four chemicals). ROIs with *I*_blank_ > 1.3 times the 99^th^ percentile of the shuffled *I*_blank_, representing neurons that putatively responded to valve and/or fluid velocity changes, were excluded. ROIs with both significant *I*_valine_ and *I*_alanine_ were defined as appetitive chemical-encoding neurons, and ROIs with both significant *I*_cadaverine_ and *I*_putrescine_ were defined as aversive chemical-encoding neurons. Valence-encoding neurons were defined as the union of appetitive and aversive chemical-encoding neurons.

### Calculation of motor and valence preference for each neuron

Each of the mutual information values were normalized by dividing by the 99^th^ percentile of the corresponding shuffled values (for simplicity of notation, hereafter referred to using the original symbols). Motor preference was defined for each motor-encoding neuron by the equation:

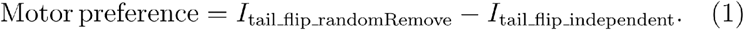

A higher value (> 0) of this quantity indicates stronger S-T event encoding, as mutual information kept after random tail flip removal (note the above-mentioned matching to S-T event proportion) is more than that with complete S-T removal. Conversely, a lower value (< 0) of such indicates stronger independent tail flip encoding, as more mutual information is kept with complete independent tail flip preservation than random removal that reduced independent tail flips while keeping some S-T events. Valence preference was defined for each sensory-encoding neuron by:

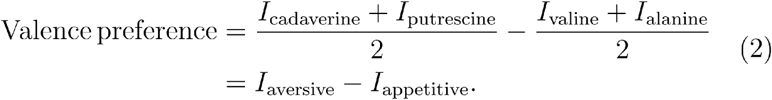

The norm of sensorimotor preferences was calculated for each unit of the union set of motor- and valence-encoding neurons that were S-T- and aversive valence-preferring (i.e., *I*_tail_flip_randomRemove_ - *I*_tail_flip_independent_ > 0 and *I*_aversive_ - *I*_appetitive_ > 0), or independent tail flip- and appetitive valence-preferring (i.e., *I*_tail_flip_randomRemove_ - *I*_tail_flip_independent_ < 0 and *I*_aversive_ - *I*_appetitive_ < 0) by the equation:

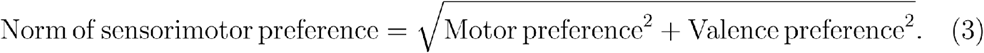

### Data visualization

All smoothing was performed using the built-in *smooth* function in MATLAB (R2018b, MathWorks, USA), with span of moving average = 3 (**Figures 2e, h, i, j, Supplementary Figures 3e, f, 4**). In the time-series stimulus-evoked plots (**Figures 3b–d**), time bins were chosen as 5, 1.67, and 5 seconds, according to the event rates of saccade (**Figure 3b**), tail flip (**Figure 3c**) and S-T events (**Figure 3d**), respectively. Each bin includes the trailing but not the leading edge. Quantities were normalized by subtracting the average binned values in the 10-second pre-stimulus windows. Tail flip frequency was normalized to and expressed as the number of standard deviations of the binned pre-stimulus frequency (**Figure 3c**).

For the mean response of motor-encoding and valence-encoding ROI (**Supplementary Figures 8a, b**), each range includes the trailing but not the leading delimiter. For the motor response trace shown (**Figures 5a, b, Supplementary Figures 8a, 9–11**), the linear trend of ΔF/F before the tail flip event (*t* = -5s to -1s, 8 or 4 imaging frames) and the ΔF/F of the first time point (at *t* = -5s) were subtracted from all data points for the visualization of motor-triggered responses.

For brain regional average analysis, regions were included only if there were data from at least 3 larvae. As habenula and caudal hypothalamus were not consistently sampled (see above anatomical imaging section), these two regions were not included in any regional average plots (**Figures 4d, f, i, k, Supplementary Figures 7b, d, f, h, j, l, 9, 11**). Other regions without neurons from sufficient larvae that passed the sensory and/or motor function identification criteria (see above neuronal function identification sections) were also not included in some of these plots.

### Statistics

In all plots, significance levels are set at *P* < 0.05. Significant *P*-values are shown in black font, while near-significant values are shown in gray font. Behavioral experiments included data from *n* = 8 larvae, comprising 838 conjugate saccades and 2,642 tail flips. The numbers of blank-control, appetitive, and aversive trials included in behavioral statistical analyses were 45, 83, and 86, respectively. For event-level analyses, repeated observations within larvae were accounted for by including larval identity as a random effect in mixed-effects models, where applicable.

For comparisons of time spent in the different quadrants in the valence determination assay (**Figure 1c**, **Supplementary Figures 1c**–**e**), two-sided *Z*-test with population standard deviation estimated from the sample standard deviation in the control group (i.e., water only in all quadrants) was used, followed by Benjamini-Hochberg procedure to control the false discovery rate across multiple comparisons. The comparisons were made across 7 cases, including mean of individual, any 2, or all 3 chemical quadrants vs. mean of quadrant I. The most significant or near-significant *P*-values for each chemical are shown.

In behavioral analyses, a two-sided Wilcoxon signed-rank test was used to compare paired larva-level median pre-event intervals in (**Figure 1e**), whereas two-sided Wilcoxon rank-sum tests were used to compare independent populations (**Supplementary Figures 4a, c**). One-sided Chi-squared test was used to compare proportions between two groups (**Supplementary Figure 4b**).

A linear mixed-effects model was used with a random intercept per larva (Delay ∼ 1 + (1|FishID)) to test whether the population-level mean delay differed from zero (fixed intercept) (**Figures 2b, c, Supplementary Figures 5a–d**). Linear mixed-effects models with event type as a fixed effect and larval identity as a random intercept were used to compare conditions (Parameter ∼ EventType + (1|FishID)) **(Figures 2d, e, i, j, Supplementary Figures 3e, f**), where Parameter denotes pre-event interval, tail flip asymmetry, or average undulation depending on the panel. For assessing expected proportions, a binomial generalized linear mixed-effects model with a random intercept per larva (Ipsilateral ∼ 1 + (1|FishID)) was used, testing whether the population-level proportion differed from 0.5 (**Figure 2f, Supplementary Figures 3b, c, 5e, f**). A binomial generalized linear mixed-effects model with larval identity as a random intercept was used to compare proportions between two groups (Laterality ∼ Case + (1|FishID)) (**Figure 2h**). Linear mixed-effects models were fitted using the MATLAB function *fitlme*, and generalized linear mixed-effects models with a binomial distribution and logit link were fitted using *fitglme* (R2024b, MathWorks, USA).

For multiple group comparisons, a linear mixed-effects model with group as a fixed effect and larval identity as a random intercept (Response ∼ Group + (1|FishID)) was used, followed by planned linear contrasts between groups. Global *P*-values for the effect of group were obtained by applying MATLAB function *anova* (R2024b, MathWorks, USA) to the fitted mixed-effects model (**Figures 3b–f)**. To probe specific differences between groups, planned linear contrasts were tested using MATLAB function *coefTest* (R2024b, MathWorks, USA) on the same model (e.g., appetitive vs aversive; **Figures 3b–f**). For **Figures 3b–d**, models were applied to data from the stimulus window and the subsequent 5-s post-stimulus period. For **Figures 3e–h** and **Supplementary Figure 6**, analyses included events occurring during the 10-s stimulus window. For comparing tail flip asymmetry and average undulation between coupled saccade–tail flip (S-T) and independent tail flip (T) events under appetitive and aversive stimulation (**Supplementary Figures 5g, h**), linear mixed-effects models were used with fixed effects of StimGroup and Subgroup and a random intercept per larva (Parameter ∼ StimGroup*Subgroup + (1|FishID)) to assess population-level effects.

To test whether the structured properties of aversive-context S-T events could arise from increased saccade occurrence alone, saccades that occurred strictly within the stimulus window (saccade onset after stimulus onset and before/coinciding with stimulus offset) were isolated for shuffling. The saccade onset and offset times were jointly circularly shifted within each aversive stimulus trial and S-T events were reidentified using the original closest-tail-flip criterion (onset difference ≤ 0.5 seconds). This preserved saccade count, duration, direction, trial identity, and the tail-flip sequence. For each of 400 shuffled datasets, the corresponding mixed-effects model used for observed aversive data was refitted with larval identity as a random intercept. One-sided empirical shuffle *P*-values were calculated as the proportion of valid shuffled datasets with a statistic at least as extreme as the observed statistic, using a +1 correction in the numerator and denominator. Tests assessed whether observed S-T events showed greater saccade-leading onset delay and ipsilateral alignment than shuffled data (**Supplementary Figures 5i, j**).

To assess the relationship between temporal coordination and tail flip kinematics (**Figures 3g, h**), linear mixed-effects models were used with Δ tail flip asymmetry or Δ average undulation as the response, PC1, PC2, group, and their interactions as fixed effects, and larval identity as a random intercept (ΔMetric ∼ PC1`*`Group + PC2`*`Group + (1|FishID)). PC1 and PC2 scores were obtained from a principal component analysis of five temporal coordination variables (**Supplementary Figure 6**). Pairwise Pearson correlations among the five temporal variables were visualized in **Supplementary Figure 6a**; correlations with unadjusted *P*-values below the Bonferroni-adjusted threshold across the 10 unique pairwise comparisons (*P* < 0.005) were displayed. Group-specific PC1 effects and the difference in PC1 slopes between appetitive and aversive groups were tested using planned contrasts on the fixed-effect coefficients (**Figures 3g, h**).

Whole-brain neuronal analyses were performed on 5 larvae, with statistical comparisons conducted at the larval level. For comparisons of motor preference, valence preference, and the norm of sensorimotor preference between the pallium and other brain regions, one-sided paired *t*-tests were performed using per-larva regional averages (**Figures 4d, f, i, k**). A two-sided paired *t*-test was used to compare the mean positive valence mutual information between the pallium and the subpallium at the larva level (**Supplementary Figure 7b**). To assess neuronal activation onset relative to motor-event onset (**Figures 5a, b, Supplementary Figures 9–11**), right-sided *Z*-tests comparing pre-chemical cue-associated event neuronal activity at the two time points preceding the motor event to zero were performed on larva-averaged neuronal traces. Significance was set at 2.5 standard deviations (S.D.), with S.D. estimated from spontaneous event baseline activity variability. Adjustment for multiple comparisons across time points and brain regions was made using the Benjamini-Hochberg procedure.

### Use of artificial intelligence tools

Perplexity (Perplexity AI, Inc.) was used for language editing and assistance with code development. All outputs were reviewed and verified by S.K.H.S., who takes full responsibility for the manuscript and its analyses.

## Data availability

Preprocessed data necessary to replicate these results are available at [https://doi.org/10.5281/zenodo.17333753].

## Code availability

The custom code for data analysis is available on GitHub [https://github.com/khsamuelsy/ChemoEyeTailMvmnts/].

## Acknowledgement

We thank Dorothy Ieong for zebrafish husbandry and technical support, and Anki Miu for administrative assistance for the project. We thank Owen Randlett for insightful discussions on the design of the project, and critical comments on the manuscript. We also thank Hui Zhao and Shen Gu for the provision of zebrafish larvae for equipment testing during initial protocol establishment. This work was supported by a Croucher Innovation Award (CIA20CU01) from the Croucher Foundation; the General Research Fund (14100122), the Collaborative Research Fund (C6027-19GF, C7074-21GF and C4062-22EF), and the Area of Excellence Scheme (AoE/M-604/16) of the RGC of UGC of Hong Kong; the Health@InnoHK program of the Innovation and Technology Commission of Hong Kong; the Excellent Young Scientists Fund (Hong Kong and Macau) (82122001) from the National Natural Science Foundation of China; the Asian Young Scientist Fellowship from the Future Science Awards Foundation; Margaret K. L. Cheung Research Centre for Management of Parkinsonism; the Lo’s Family Charity Fund Limited; Gerald Choa Neuroscience Institute Research Program Fund.

## Author contributions

Conceptualization: S.K.H.S., H.K.

Methodology: S.K.H.S., C.F., K.W., Y.M., Y.H., H.K.

Investigation: S.K.H.S., D.C.W.C., J.J.Z., J.L., C.F., Y.H., H.K.

Data Curation: S.K.H.S.

Formal Analysis: S.K.H.S.

Software: S.K.H.S., H.K.

Validation: S.K.H.S., H.K.

Visualization: S.K.H.S.

Supervision: H.K.

Project Administration: S.K.H.S., V.C.T.M., K.K.Y.W., H.K.

Funding Acquisition: V.C.T.M., K.K.Y.W., H.K.

Writing - Original Draft: S.K.H.S., D.C.W.C., H.K.

Writing - Review & Editing: All authors.

---

**Supplementary Figure 1.**
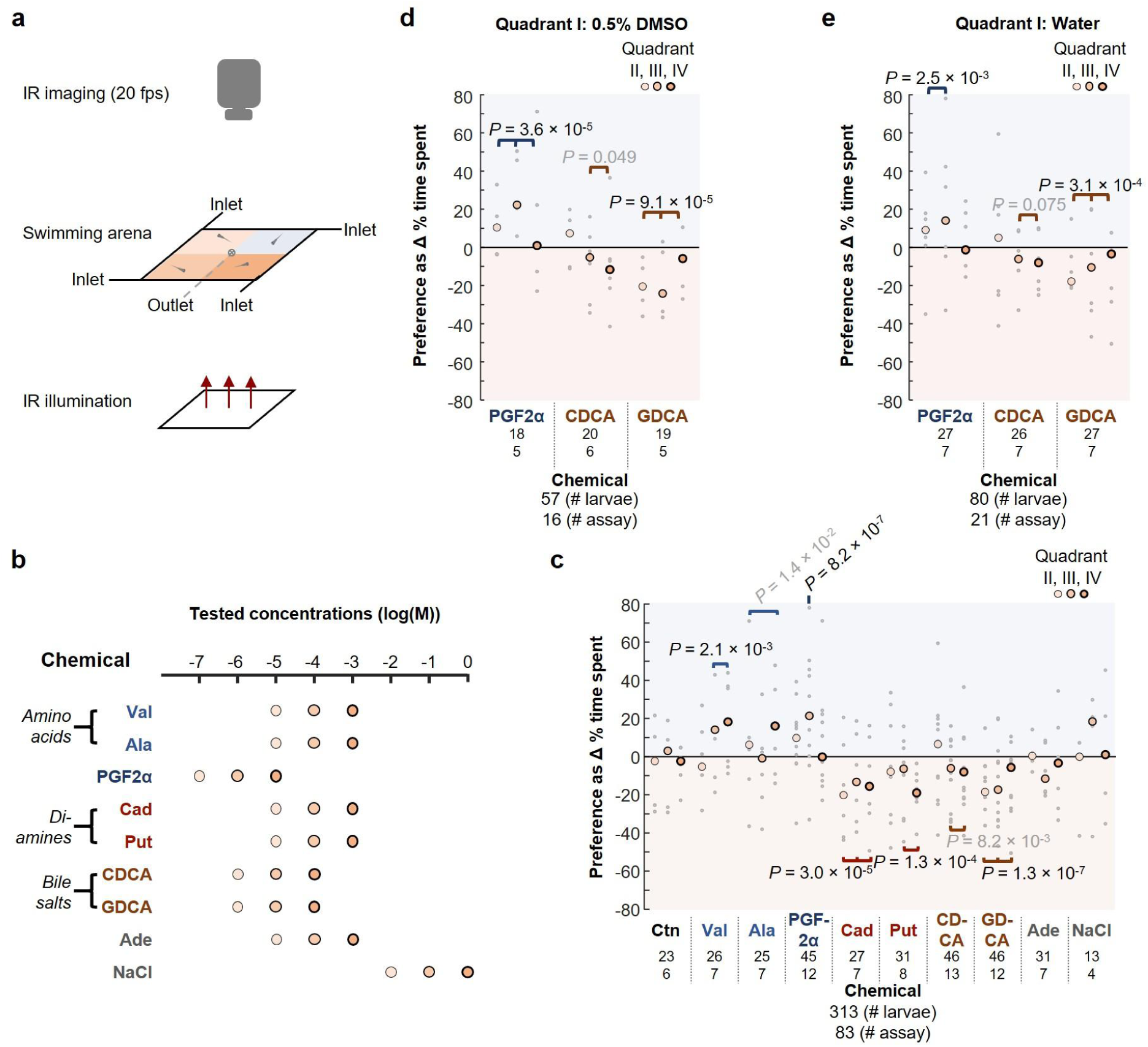
Schematics and additional analysis of the valence determination assay. **(a)** Schematic of the valence determination assay. Zebrafish larvae swimming in a two-dimensional arena (40 mm × 40 mm × 1.5 mm) are imaged at 20 fps under infrared (IR) illumination in the absence of visible light. Four quadrants with equal area are created and maintained by a constant, slow inflow of fluid at each corner, with outflow at a shared central outlet. The laminar flow maintains static borders between the zones. **(b)** Assayed chemicals and tested concentrations (varied across 3 consecutive orders of magnitude for each), including valine (Val), alanine (Ala), prostaglandin F2α (PGF2α), cadaverine (Cad), putrescine (Put), chenodeoxycholic acid (CDCA), glycodeoxycholic acid (GDCA), adenosine (Ade), and sodium chloride (NaCl). These were dissolved either in water or 0.5% DMSO and infused into quadrant II (lowest concentration) to IV (highest concentration). **(c)** Differences in % time spent in the chemical quadrants (i.e., II–IV, with different and increasing concentrations for each of the tested chemicals as indicated in **(b)**) compared with the control quadrant (i.e., I), showing the values of individual assays (gray dots) and the medians (black-outlined dots). **(d)** & **(e)** Differences in % time spent in the chemical quadrants (i.e., II**–**IV, with different and increasing concentrations for each of the tested chemicals as indicated in **(b)**) compared with the water quadrant (i.e., I), showing the values of individual assays (gray dots) and the medians (black-outlined dots). The experiments were performed with either **(d)** 0.5% DMSO or **(e)** water in quadrant I. In **(c)**–**(e)**, below each chemical name, the first number indicates the total number of larvae assayed, and the second number specifies the total number of assays performed. *P*-values: For each chemical, multiple *Z*-tests comparing mean differences in % time between the control quadrant I and all individual non-control quadrants and their possible combinations, against a null hypothesis of zero difference. The displayed *P* value is from the most significant comparison. The population standard deviation (S.D.) was estimated from the sample S.D. of the control group (see **Methods**). Statistical significance was determined using a Benjamini–Hochberg-derived significance threshold applied to the unadjusted *P*-values (controlled within each odor). Black, significant; gray, near-significant.

**Supplementary Figure 2.**
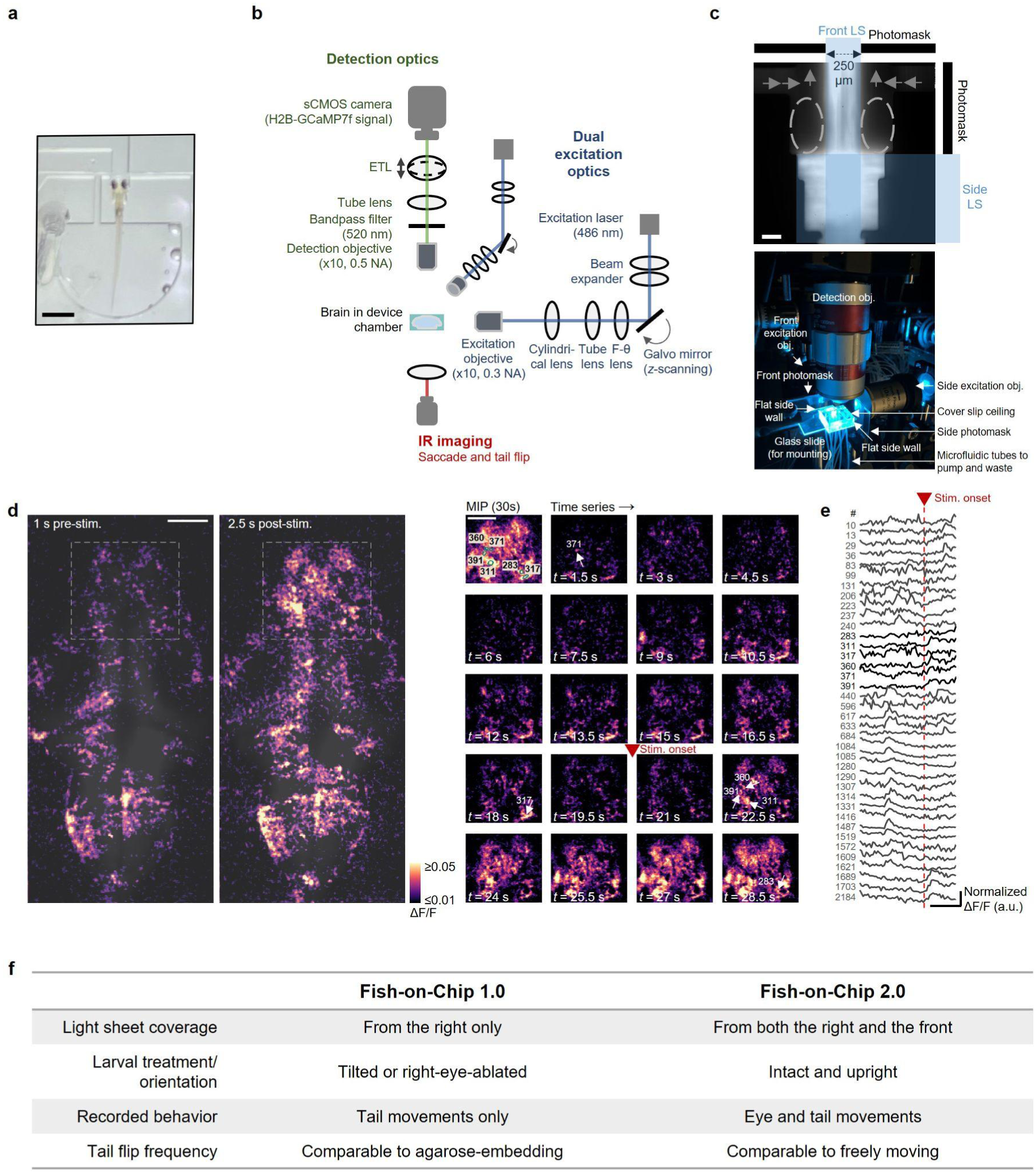
More details on Fish-on-Chip 2.0: Enhanced platform for naturalistic behavioral-neural readouts in tethered larval zebrafish. **(a)** A photograph of a tethered larval zebrafish loaded into a PDMS-based microfluidic device (scale bar: 1 mm). Its eyes and tail are free to move while the head is fixed in place for fluorescence imaging. **(b)** Optical layout of the light sheet microscope, including two light sheet scanning modules which scan the larval brain from the side and the front respectively, a fluorescence detection module, and an IR imaging module for saccade and tail movements recording. **(c)** Upper panel: A fluorescence image showing sodium fluorescein in the microfluidic device excited by the front and side scanning light sheets (LS, illustrated by overlaid blue shadows), with photomasks placed to avoid the light sheets from directly reaching the larval eye positions (marked with white dashed ovals). A 250-µm window in the front photomask allows for the passage of the front excitation light to scan the brain regions between the eyes. Scale bar: 100 µm. Lower panel: A photograph showing the microfluidic device, objectives, and the associated tubings. **(d)** Left panel: An example functional image plane showing neural activation at two time points: before and after the presentation of 1 mM cadaverine. Anatomical reference images are overlaid. Right panel: A maximum intensity projection (MIP) and the time-series images of the first 30-second interval, focusing on a zoomed-in forebrain region indicated (dashed white rectangle on the left). 6 regions-of-interest (ROIs) are outlined. The red arrow marks cadaverine stimulus onset. White arrows indicate the locations and times of each ROIs near a calcium event’s peak. Scale bars: 100 μm. **(e)** The corresponding calcium signal traces of 38 representative neurons recorded before and after the presentation of 1 mM cadaverine, with onset indicated by the red arrow and dotted red line. Scale bar: 10 seconds. **(f)** Summary of key differences between Fish-on-Chip 1.0 and 2.0. The enhanced 2.0 platform eliminates the need for tilting or right-eye ablation in larvae, allowing for naturalistic, whole-body and brain-wide behavioral-neural recordings during spontaneous behavior and chemosensory cue presentations.

**Supplementary Figure 3.**
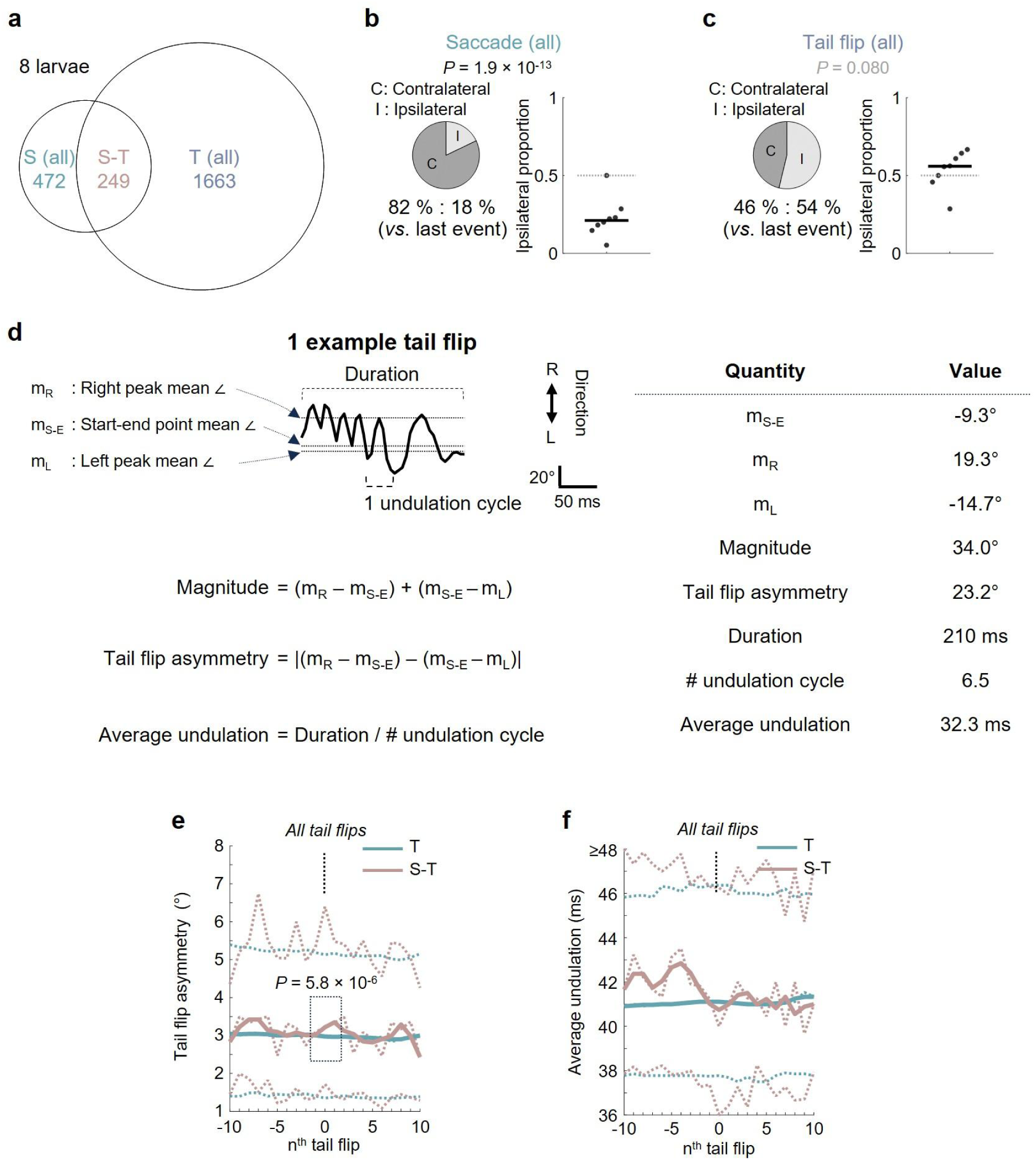
Additional analysis of spontaneous saccade-tail flip coordination. **(a)** Venn diagram showing all counts of spontaneous saccades, tail flips and S-T events (*n* = 8 larvae). **(b)** & **(c)** Left panels: Pie charts showing the proportions of contra- and ipsilateral turns, comparing the directionality of **(b)** saccades and their preceding events, and **(c)** tail flips and their preceding events. Right panels: Corresponding larva-based ipsilateral proportion. The horizontal lines indicate the corresponding medians across larvae. *P*-values: Binomial generalized linear mixed-effects model with a random intercept per larva (ipsilateral vs. contralateral), testing whether the ipsilateral fraction differs from 0.5. **(d)** Illustration of the various kinematic parameters of an example tail flip. Dashed lines indicate the right peak mean angle, start-end point mean angle, and left peak mean angle. **(e)** Tail flip asymmetry and **(f)** average undulation of self and flanking tail flips for all S-T events or all independent tail flips (T). The thin dotted curves represent the medians, 25^th^, and 75^th^ percentiles of all events, and thick solid lines show the corresponding moving-average curves of the medians. *P*-value: Linear mixed-effects model with event type as a fixed effect and larval identity as a random intercept, applied to events pooled across the peak lag and its adjacent ±1 lags. Data were collected from *n* = 8 larvae, with 472 spontaneous saccades and 1,663 spontaneous tail flips.

**Supplementary Figure 4.**
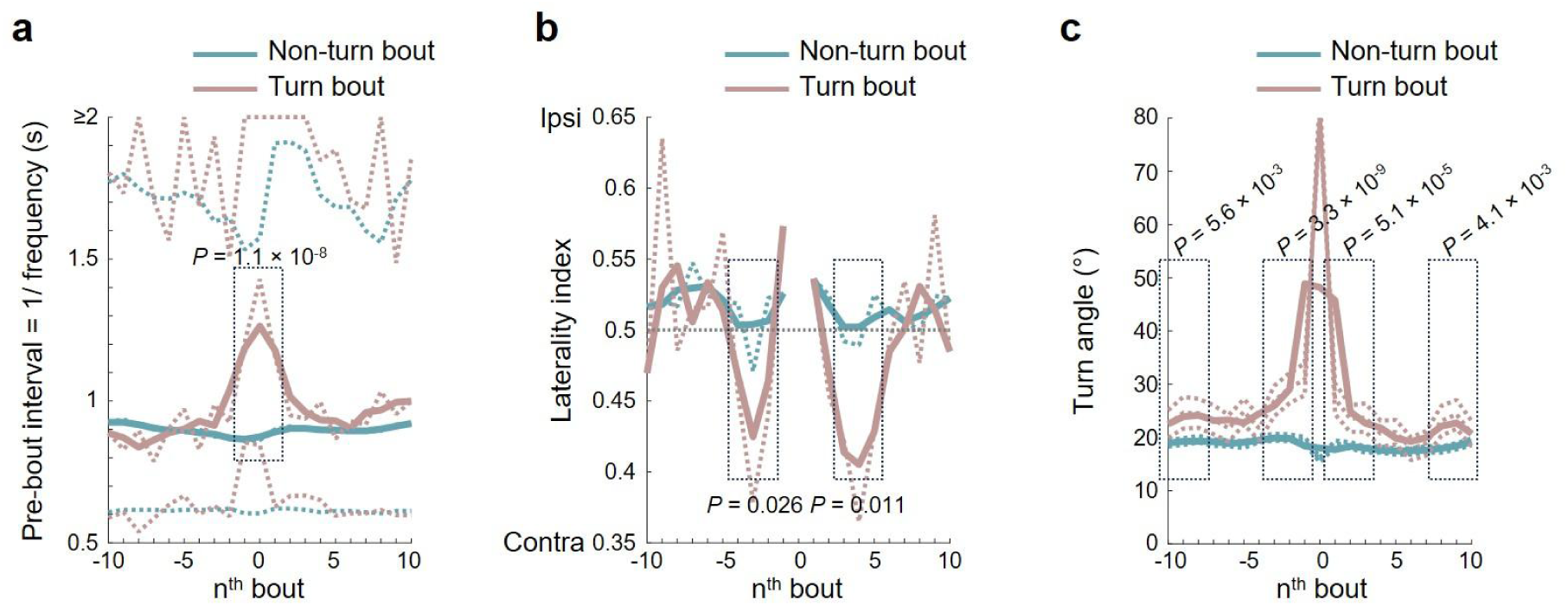
Freely moving zebrafish larvae dynamically align bouts in rate and direction for spontaneous body turns. **(a)** Pre-event interval [1/ frequency] of self and flanking swim bouts for non-turn (teal) or turn (coral) swim bouts (see **Methods**). Thin dotted lines represent the raw median values, 25^th^, and 75^th^ percentiles. Thick solid lines are the moving-average of medians. **(b)** Laterality index (defined as proportion of ipsilateral turn) in flanking swim bouts for non-turn (teal) or turn (coral) events. The raw median values (thin dotted lines) and corresponding moving-average curves (thick solid lines) are shown. *P*-values: One-sided Chi-squared test comparing peaks and ± 1 events. **(c)** Turn angle of self and flanking tail flips for non-turn (teal) or turn (coral) swim bouts. The thin dotted lines represent mean ± S.E.M. across time, and thick solid lines show the corresponding moving-average curves. In **(a)** & **(c)**, *P*-values: Two-sided Wilcoxon rank-sum test comparing peaks and ± 1 events. Note that however in **(c)**, the centered bout (i.e., 0^th^ next bout) is by definition different and therefore only flanking bouts are compared. For all plots, data were collected from *n* = 11 larvae, with 1,529 spontaneous swim bouts.

**Supplementary Figure 5.**
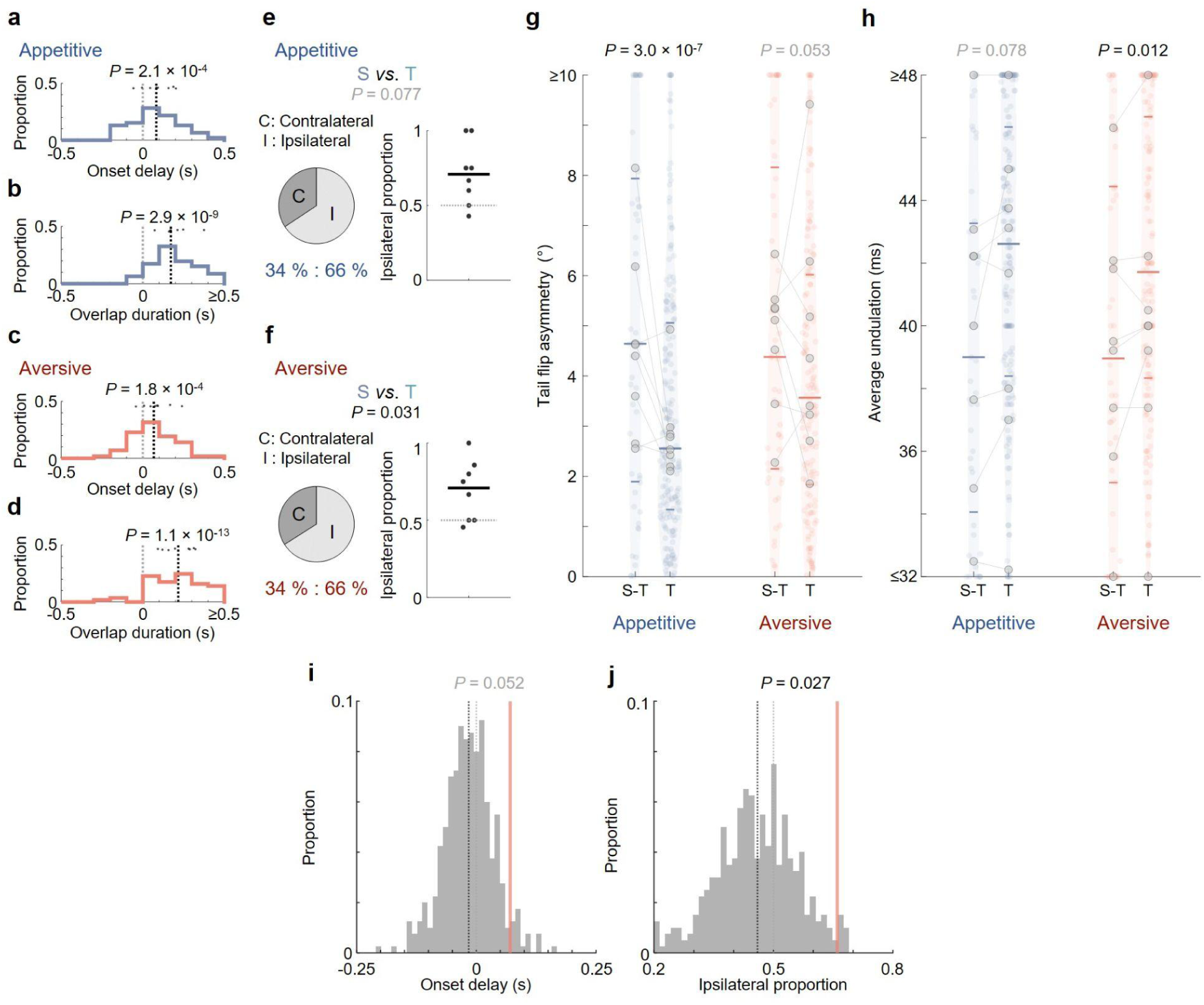
Conserved temporal, directional and kinematic characteristics of coupled saccade-tail flip (S-T) events under chemical stimulus presentation. **(a)–(d)** Histograms of saccade-tail flip (a) & (c) onset delay, and (b) & (d) overlap duration for S-T events during (a) & (b) appetitive and (c) & (d) aversive chemical stimulus presentation. Note that the precision of measurements for the onset and offset for saccade and tail flip were 0.0625s and 0.0025s due to the respective sampling rates. Black and gray dashed lines indicate overall medians and zero, respectively. Dots indicate the medians for each larva. *P*-values: Linear mixed-effects model with a random intercept per larva, testing the fixed intercept against zero. **(e)**–**(f)** Left panels: Pie charts showing the proportions of contra- and ipsilateral turns, comparing the directionality of saccades and turns in S-T events during **(e)** appetitive and **(f)** aversive chemical stimulus presentation. Right panels: Corresponding larva-based ipsilateral proportion. *P*-values: Binomial generalized linear mixed-effects model with a random intercept per larva (ipsilateral vs. contralateral), testing whether the ipsilateral fraction differs from 0.5. **(g,h) (g)** Tail flip asymmetry and **(h)** average undulation (for tail flips with ≥ 4 undulation cycles) of S-T events and independent tail flips (T) during appetitive and aversive chemical stimulus presentation. Horizontal lines indicate medians, 25^th^, and 75^th^ percentiles. Shadows of the violin plots scale according to the probability density function. Gray dots indicate the medians for each larva in each condition, and gray lines connect the same larva across S-T and T, illustrating within-animal changes. *P*-values: S-T vs. T planned contrasts within each stimulus group, derived from a linear mixed-effects model with fixed effects of StimGroup, Subgroup, and their interaction, and larval identity as a random intercept. **(i,j)** Aversive-trial within-trial circular saccade-time shuffle controls for **(i)** onset delay and **(j)** ipsilateral alignment. Saccade times were jointly shifted within trial and S-T events reidentified using the original ≤ 0.5 seconds nearest-tail-flip criterion. Gray histograms show 400 shuffle-derived statistics (see **Methods**); red solid and black dashed lines indicate observed statistics and median shuffled statistics, respectively. Gray dashed lines indicate zero and chance level (i.e., 0.5) respectively. One-sided empirical shuffle *P*-values are shown.

**Supplementary Figure 6.**
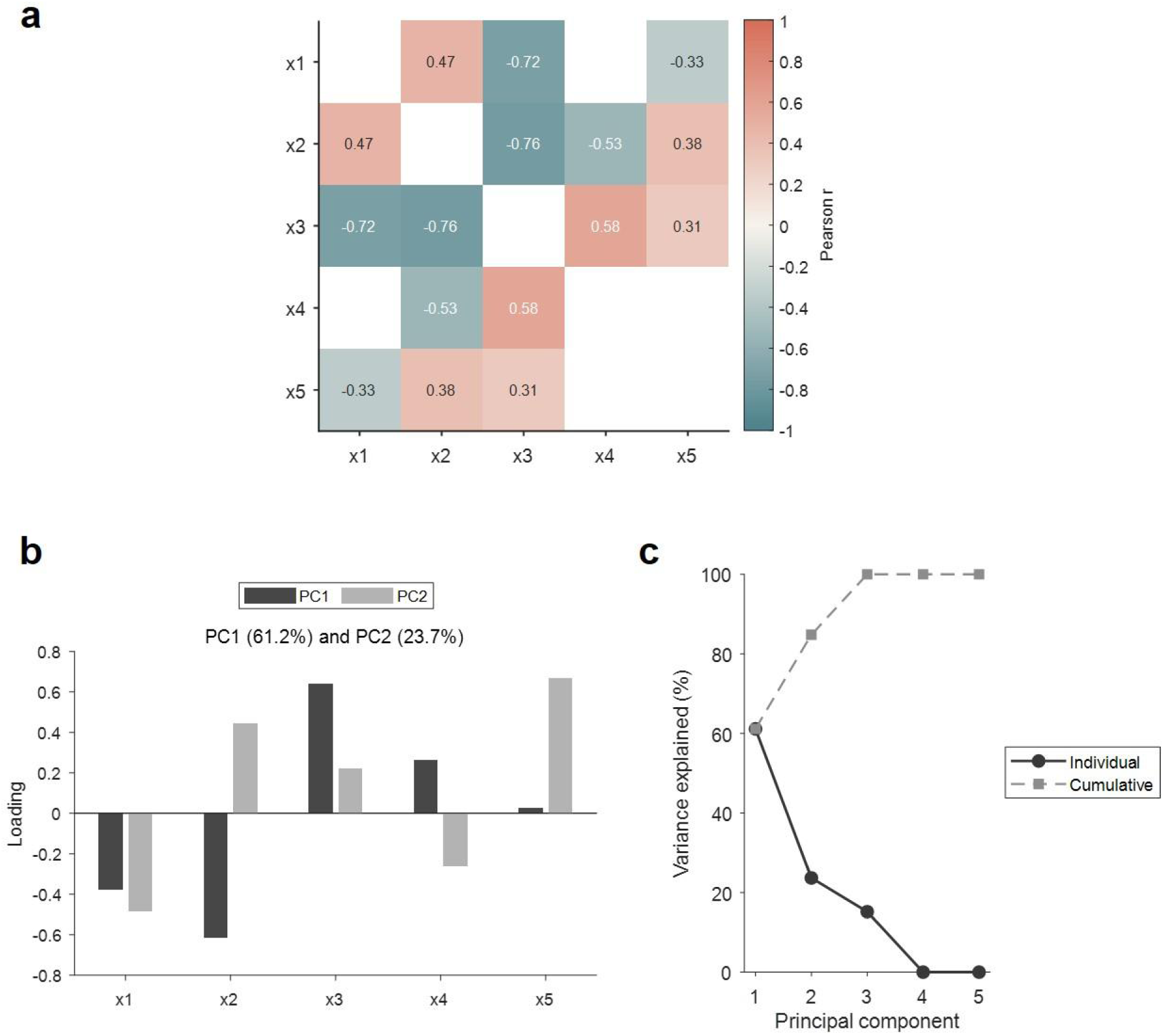
Principal component analysis (PCA) of temporal parameters in coupled saccade-tail flip (S-T) events during chemical stimulus presentation. **(a)** Pearson correlation matrix for the five temporal predictors *x*1–*x*5 describing S-T event structure, *x*1 (onset delay: tail flip onset-saccade onset), *x*2 (offset delay: tail flip offset-saccade offset), *x*3 (overlap duration: saccade offset-tail flip onset), *x*4 (saccade duration), *x*5 (tail flip duration), pooled across all trials and stimulus conditions. The color scale encodes Pearson’s *r* from −1 to 1 and numeric values indicate the exact correlations, showing that several predictor pairs are highly and statistically significantly correlated (using the Bonferroni-derived threshold). (b) Loadings of the temporal predictors (onset delay, offset delay, overlap duration, saccade duration, tail flip duration) on the first two principal components (PC1, PC2) computed from the pooled dataset. Positive and negative loadings indicate the direction and strength with which each temporal feature contributes to PC1 and PC2, with the first two components together capturing most of the variance in the temporal structure. (c) Scree plot of the PCA on the temporal predictors, showing the proportion of variance explained by each principal component (solid line, circles) and the cumulative variance explained (dashed line, squares). The first two components account for the majority of the total variance, supporting the use of PC1 and PC2 as low-dimensional summaries of the temporal structure in the mixed-effects model in Figures 3g, h.

**Supplementary Figure 7.**
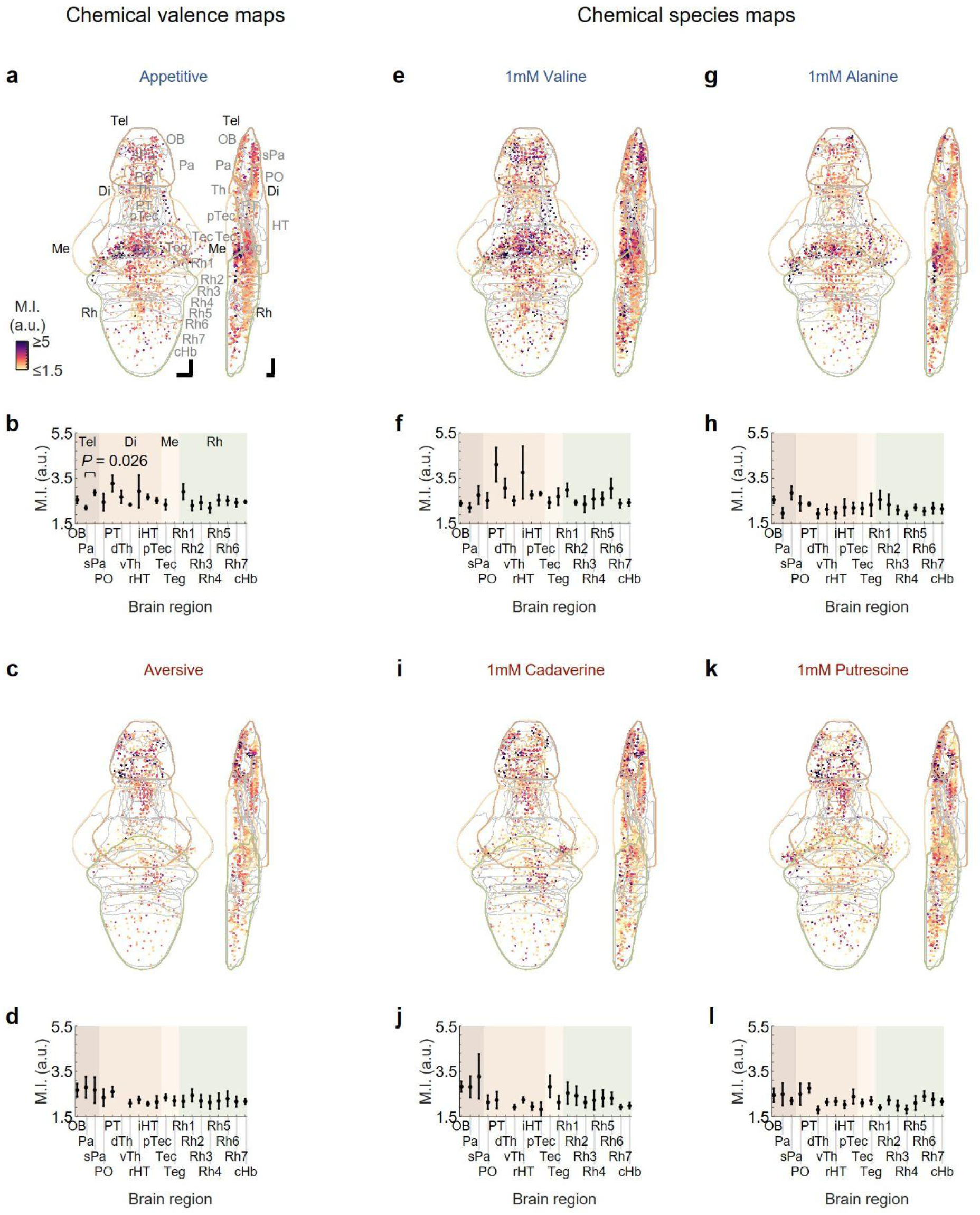
Brain-wide chemical representation. **(a)** Mean intensity (to transverse and sagittal planes) of mean mutual information (M.I.) between the calcium signals of appetitive valence-encoding regions-of-interest (ROIs) and the appetitive stimulus regressors ((*I*_valine_ + *I*_alanine_)/2, see **Methods**). **(b)** Regional mean ± S.E.M. (across larvae) of mean appetitive M.I. ((*I*_valine_ + *I*_alanine_)/2), averaged across ROIs within each region and larvae. *P*-value: Two-sided paired *t*-test comparing the pallium and the subpallium at the larval level (*n* = 5 larvae). **(c)** & **(d)** Similar to **(a)** & **(b)** but for mean aversive chemical M.I. ((*I*_cadaverine_ + *I*_putrescine_)/2) and aversive valence-encoding ROIs. **(e)** Mean intensity projections (to transverse and sagittal planes) of M.I. between the calcium signals of 1 mM valine-encoding ROIs and the stimulus regressor (*I*_valine_, see **Methods**). **(f)** Regional mean ± S.E.M. (across larvae) of sensory M.I, averaged across ROIs within each region and larvae. **(g)**–**(l)** Similar to **(e)** & **(f)** but for **(g)** & **(h)** alanine (*I*_alanine_), **(i)** & **(j)** cadaverine (*I*_cadaverine_) and **(k)** & **(l)** putrescine (*I*_putrescine_), all at 1 mM. The major brain regions (Tel: telencephalon, Di: diencephalon, Me: mesencephalon and Rh: rhombencephalon) are color-coded. Abbreviations of brain regions: Same as in Figure 4d. In all plots, the neural activity data are from *n* = 5 larvae and correspond to a subset of the behavioral dataset shown in Figures 2 & **3** (*n* = 8). Scale bars: 50 µm in the Z-Brain Atlas space.

**Supplementary Figure 8.**
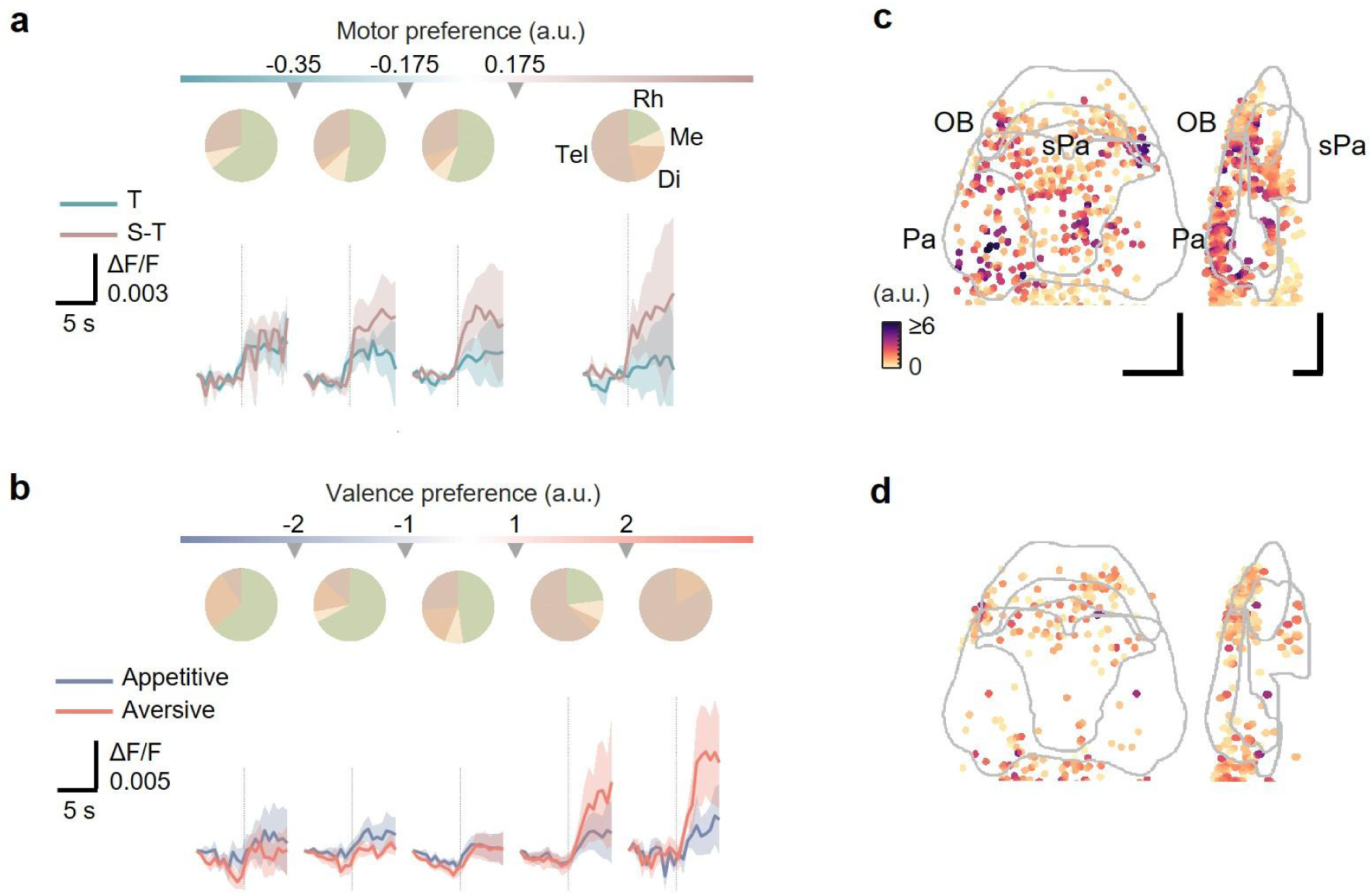
Additional analysis of brain-wide valence and motor representations. **(a)** Upper panel: Pie charts showing the regional distributions of regions-of-interest (ROIs) (median proportions across larvae) with motor preference in different ranges. Lower panel: Mean ± S.E.M. (across larvae, using per-larva averages) responses of motor-encoding ROIs with motor preference in different ranges. Dotted vertical lines indicate the times of tail flip onsets (in a corresponding neuronal imaging frame). **(b)** Upper panel: Pie charts showing the regional distributions of ROIs (median proportions across larvae) with valence preference in different ranges. Lower panel: Mean ± S.E.M. (across larvae, using per-larva averages) responses of valence-encoding ROIs with valence preference in different ranges. Dotted vertical lines indicate the time points of stimulus onsets (between two neuronal imaging frames). In **(a)** and **(b)**, the major brain regions (Tel: telencephalon, Di: diencephalon, Me: mesencephalon and Rh: rhombencephalon) are color-coded. **(c)** & **(d)** Zoom-in maps showing the forebrain regions olfactory bulb, pallium and subpallium ROIs preferring **(c)** S-T event and aversive valence in Figure 4h, and **(d)** independent tail flip and appetitive valence in Figure 4j. Scale bars: 50 μm in the Z-Brain Atlas space. Motor-preference ranges in **(a)**, from left to right, include 305, 148, 1607, and 208 motor-encoding ROIs pooled across the 5 larvae, and valence-preference ranges in **(b)**, from left to right, include 275, 592, 1199, 227, and 97 valence-encoding ROIs pooled across the 5 larvae. The neural activity data are from *n* = 5 larvae and correspond to a subset of the behavioral dataset shown in Figures 2 & **3** (*n* = 8).

**Supplementary Figure 9.**
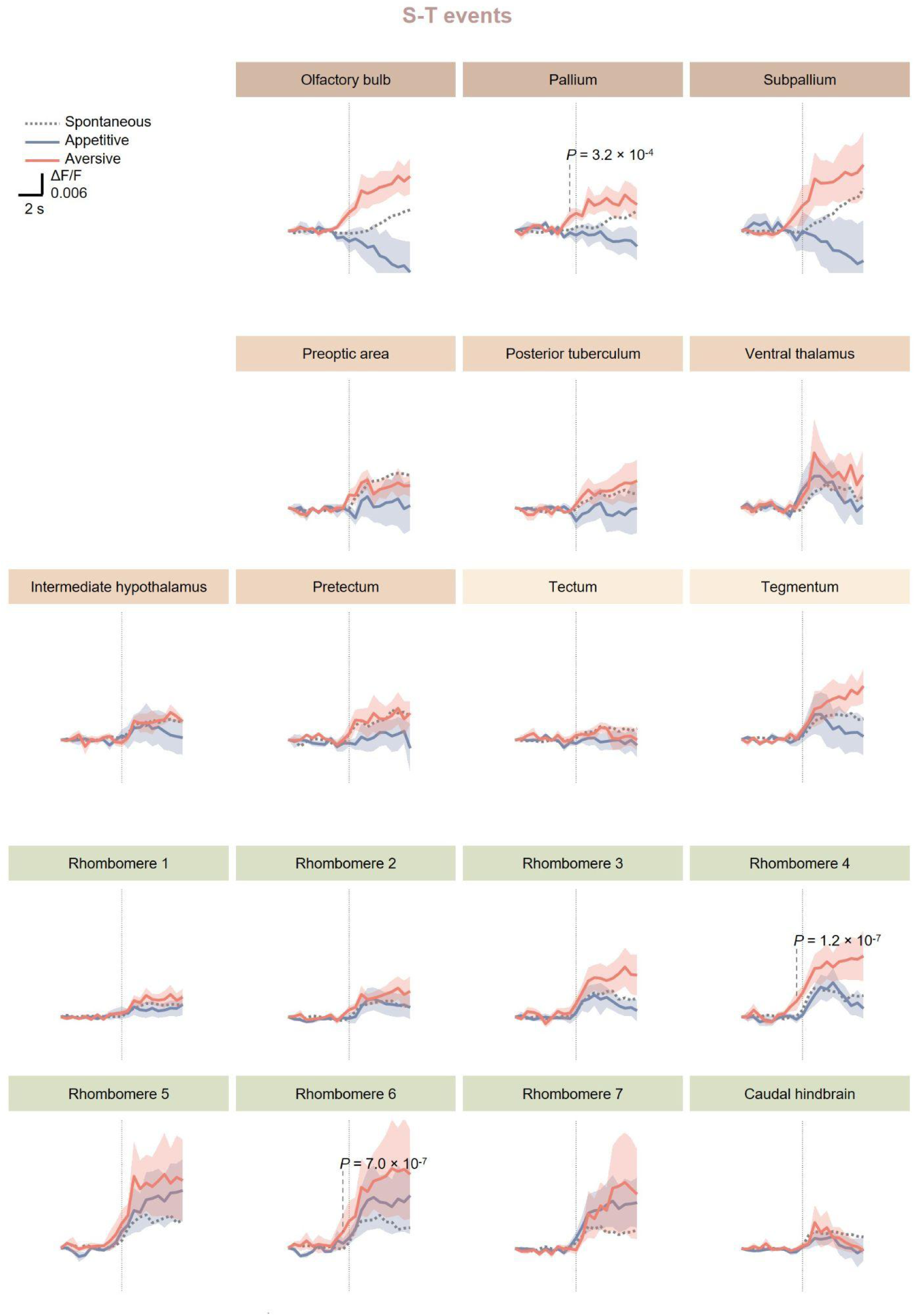
Neuronal activity associated with chemosensory-driven coupled saccade-tail flip (S-T) events across different brain regions. Mean ± S.E.M. (across *n* = 5 larvae, using per-larva averages) responses of motor-encoding regions-of-interest (ROIs) in the different brain regions, aligned to the onset of chemosensory cue-associated S-T events. Dotted traces show the corresponding neuronal responses of spontaneous S-T events. Dotted vertical lines indicate tail flip onsets (in a corresponding neuronal imaging frame). *P*-values: Right-sided *Z*-tests comparing pre-chemical cue-associated event neuronal activity at each of the two time points preceding the motor event to zero at the larval level (*n* = 5 larvae). Significance was set at 2.5 standard deviations (S.D.), with S.D. estimated from spontaneous event baseline activity variability. Adjustment for multiple comparisons across time points and brain regions was made using the Benjamini-Hochberg procedure. The neural activity data are from *n* = 5 larvae and correspond to a subset of the behavioral dataset shown in Figures 2 & **3** (*n* = 8).

**Supplementary Figure 10.**
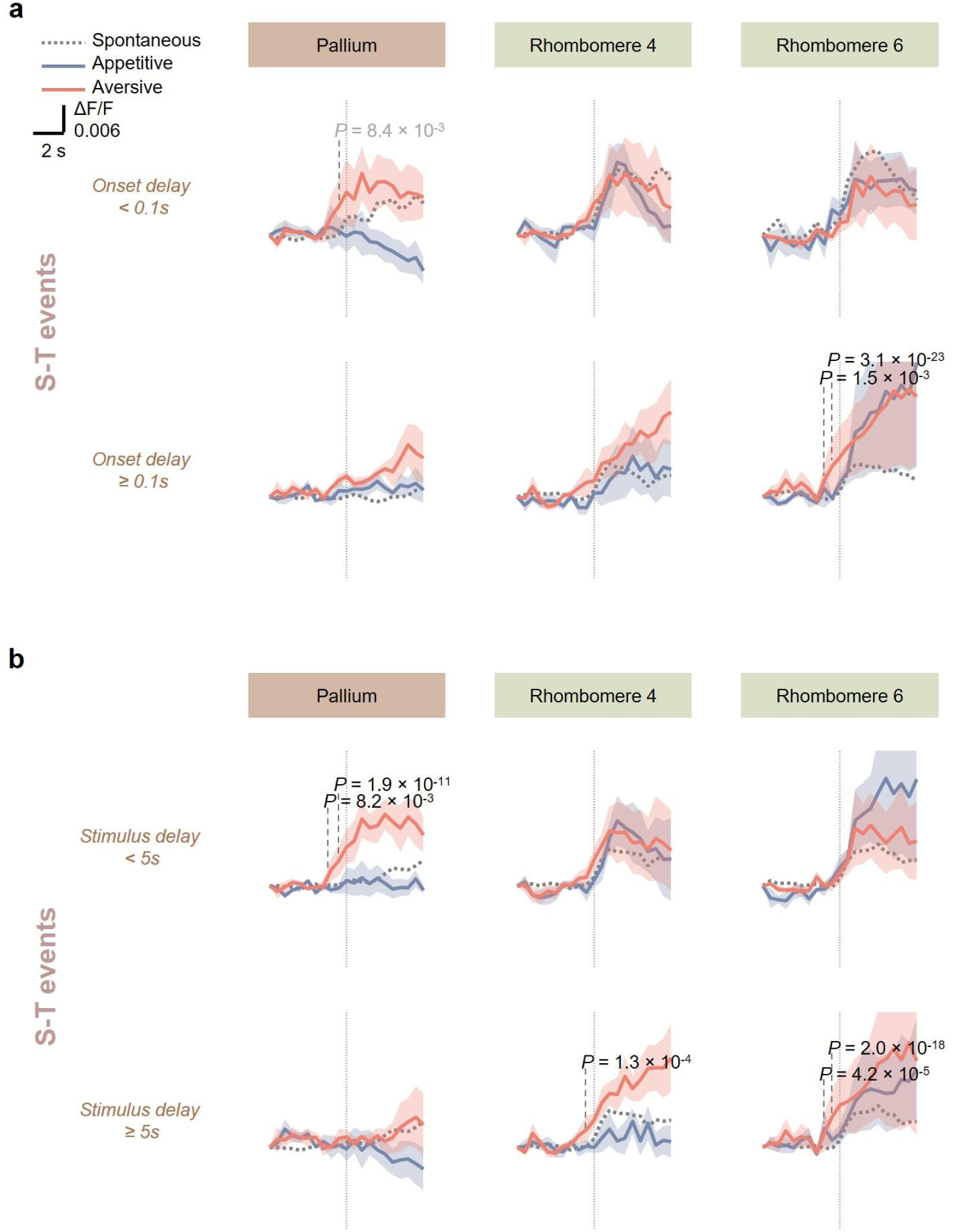
Additional analysis of neural activity underlying chemosensory-driven coupled saccade-tail flip (S-T) events. Mean ± S.E.M. (across *n* = 5 larvae, using per-larva averages) responses of motor-encoding regions-of-interest (ROIs) in the pallium, rhombomere 4 and rhombomere 6, classified according to **(a)** saccade-tail flip onset delay (tail flip onset-saccade onset) [< 0.1 seconds (upper) or ≥ 0.1 seconds (lower)], as defined in Figure 2a, and **(b)** stimulus-tail flip delay (tail flip onset-stimulus onset) [< 5 seconds (upper) or ≥ 5 seconds (lower)], and aligned to the onset of chemosensory cue-associated S-T events. Dotted traces show the corresponding neuronal responses of spontaneous S-T events (note that in **(b)**, spontaneous S-T events were not classified according to stimulus-tail flip delay). Dotted vertical lines indicate tail flip onsets (in a corresponding neuronal imaging frame). *P*-values: Right-sided *Z*-tests comparing pre-chemical cue-associated event neuronal activity at each of the two time points preceding the motor event to zero at the larval level (*n* = 5 larvae). Significance was set at 2.5 standard deviations (S.D.), with S.D. estimated from spontaneous event baseline activity variability. Adjustment for multiple comparisons across time points and brain regions was made using the Benjamini-Hochberg procedure. Statistical significance: Black, significant; gray, near-significant. The neural activity data are from *n* = 5 larvae and correspond to a subset of the behavioral dataset shown in Figures 2 & **3** (*n* = 8).

**Supplementary Figure 11.**
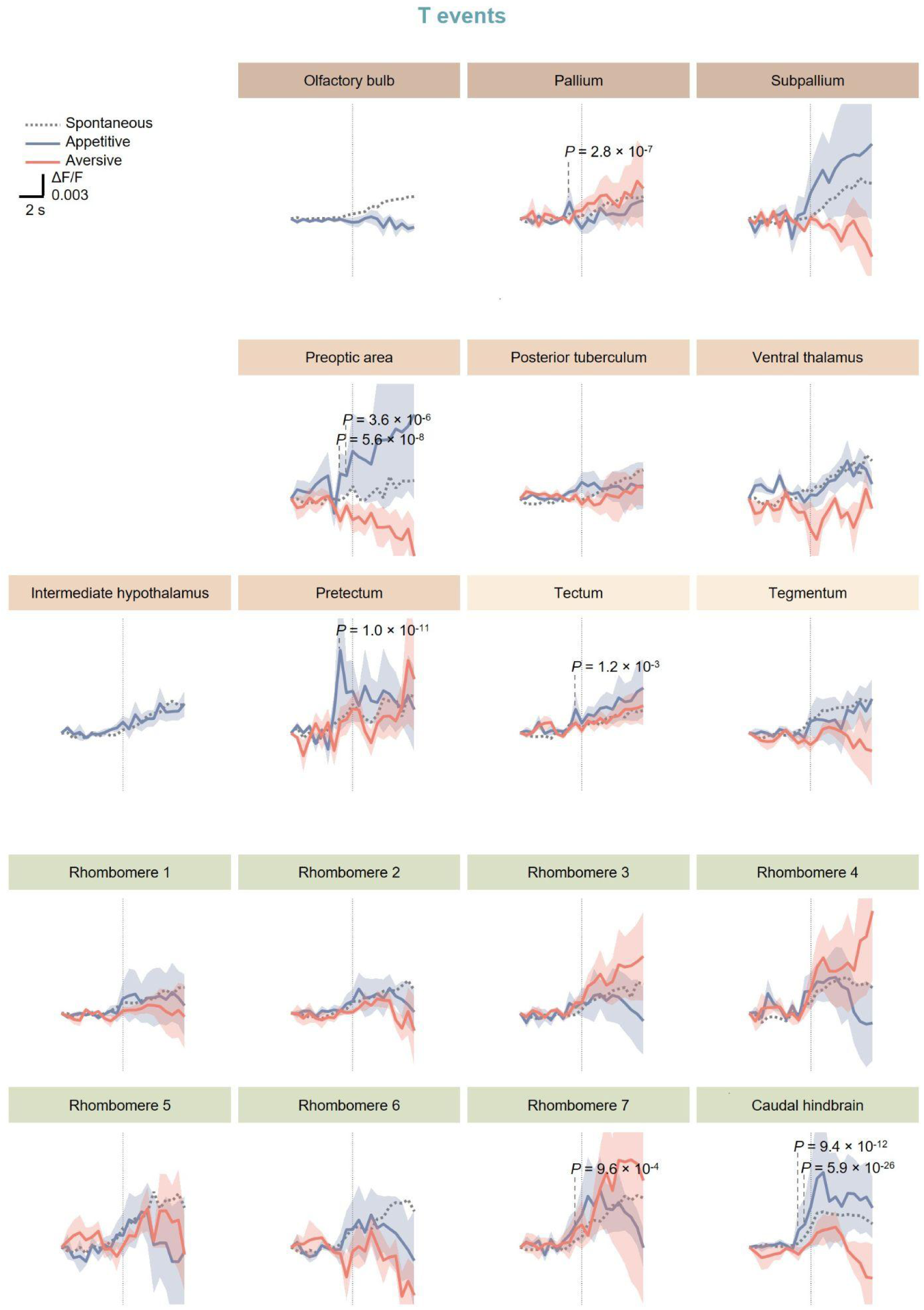
Neuronal activity associated with chemosensory-driven independent tail flips (T events) across brain regions. Mean ± S.E.M. (across *n* = 5 larvae, using per-larva averages) responses of motor-encoding regions-of-interest (ROIs) in the different brain regions, aligned to the onset of independent tail flips during chemical cue presentation. Dotted traces show the corresponding neuronal responses of spontaneous independent tail flips. Dotted vertical lines indicate tail flip onsets (in a corresponding neuronal imaging frame). *P*-values: Right-sided *Z*-tests comparing pre-chemical cue-associated event neuronal activity at each of the two time points preceding the motor event to zero at the larval level (*n* = 5 larvae). Significance was set at 2.5 standard deviations (S.D.), with S.D. estimated from spontaneous event baseline activity variability. Adjustment for multiple comparisons across time points and brain regions was made using the Benjamini-Hochberg procedure. The neural activity data are from *n* = 5 larvae and correspond to a subset of the behavioral dataset shown in Figures 2 & **3** (*n* = 8).

## References

1. Dakin, R., Fellows, T. K. & Altshuler, D. L. Visual guidance of forward flight in hummingbirds reveals control based on image features instead of pattern velocity. Proc Natl Acad Sci USA 113, 8849–8854 (2016).

2. Byrne, R. A., Kuba, M. J., Meisel, D. V., Griebel, U. & Mather, J. A. Octopus arm choice is strongly influenced by eye use. Behav. Brain Res. 172, 195–201 (2006).

3. Bianco, I. H. & Engert, F. Visuomotor transformations underlying hunting behavior in zebrafish. Curr. Biol. 25, 831–846 (2015).

4. Land, M. F. Motion and vision: why animals move their eyes. J. Comp. Physiol. A 185, 341–352 (1999).

5. Srivastava, A., Ahmad, O. F., Pacia, C. P., Hallett, M. & Lungu, C. The Relationship between Saccades and Locomotion. J. Mov. Disord. 11, 93–106 (2018).

6. Wolf, S. et al. Sensorimotor computation underlying phototaxis in zebrafish. Nat. Commun. 8, 651 (2017).

7. Dowell, C. K., Lau, J. Y. N., Antinucci, P. & Bianco, I. H. Kinematically distinct saccades are used in a context-dependent manner by larval zebrafish. Curr. Biol. (2024) doi:10.1016/j.cub.2024.08.008.

8. Imai, T., Moore, S. T., Raphan, T. & Cohen, B. Interaction of the body, head, and eyes during walking and turning. Exp. Brain Res. 136, 1–18 (2001).

9. Lohnes, C. A. & Earhart, G. M. Saccadic eye movements are related to turning performance in Parkinson disease. J Parkinsons Dis 1, 109–118 (2011).

10. Lambert, F. M., Combes, D., Simmers, J. & Straka, H. Gaze stabilization by efference copy signaling without sensory feedback during vertebrate locomotion. Curr. Biol. 22, 1649–1658 (2012).

11. França de Barros, F., et al. Conservation of locomotion-induced oculomotor activity through evolution in mammals. Curr. Biol. 32, 453–461.e4 (2022).

12. Bacqué-Cazenave, J., Courtand, G., Beraneck, M., Lambert, F. M. & Combes, D. Temporal Relationship of Ocular and Tail Segmental Movements Underlying Locomotor-Induced Gaze Stabilization During Undulatory Swimming in Larval Xenopus. Front. Neural Circuits 12, 95 (2018).

13. Lavian, H., Prat, O., Petrucco, L., Štih, V. & Portugues, R. The representation of visual motion and landmark position aligns with heading direction in the zebrafish interpeduncular nucleus. BioRxiv (2024) doi:10.1101/2024.09.25.614953.

14. Sugioka, T., Tanimoto, M. & Higashijima, S.-I. Biomechanics and neural circuits for vestibular-induced fine postural control in larval zebrafish. Nat. Commun. 14, 1217 (2023).

15. Ehrlich, D. E. & Schoppik, D. A primal role for the vestibular sense in the development of coordinated locomotion. eLife 8, (2019).

16. Semmelhack, J. L. et al. A dedicated visual pathway for prey detection in larval zebrafish. eLife 3, (2014).

17. Mearns, D. S., Donovan, J. C., Fernandes, A. M., Semmelhack, J. L. & Baier, H. Deconstructing Hunting Behavior Reveals a Tightly Coupled Stimulus-Response Loop. Curr. Biol. 30, 54–69.e9 (2020).

18. Ahrens, M. B. et al. Brain-wide neuronal dynamics during motor adaptation in zebrafish. Nature 485, 471–477 (2012).

19. Naumann, E. A. et al. From Whole-Brain Data to Functional Circuit Models: The Zebrafish Optomotor Response. Cell 167, 947–960.e20 (2016).

20. Baden, T. Ancestral photoreceptor diversity as the basis of visual behaviour. *Nat*. Ecol. Evol. 8, 374–386 (2024).

21. Currier, T. A. & Nagel, K. I. Multisensory control of navigation in the fruit fly. Curr. Opin. Neurobiol. 64, 10–16 (2020).

22. Rössler, W. Multisensory navigation and neuronal plasticity in desert ants. Trends Neurosci. 46, 415–417 (2023).

23. Danilovich, S. & Yovel, Y. Integrating vision and echolocation for navigation and perception in bats. Sci. Adv. 5, eaaw6503 (2019).

24. Radvansky, B. A., Oh, J. Y., Climer, J. R. & Dombeck, D. A. Behavior determines the hippocampal spatial mapping of a multisensory environment. Cell Rep. 36, 109444 (2021).

25. Davis, S. N., Zhu, Y. & Schoppik, D. Larval zebrafish maintain elevation with multisensory control of posture and locomotion. J. Exp. Biol. (2025) doi:10.1242/jeb.250087.

26. Hoover, K. C. Smell with inspiration: the evolutionary significance of olfaction. Am. J. Phys. Anthropol. 143 **Suppl 51**, 63–74 (2010).

27. Sy, S. K. H. et al. An optofluidic platform for interrogating chemosensory behavior and brainwide neural representation in larval zebrafish. Nat. Commun. 14, 227 (2023).

28. Wakisaka, N. et al. An adenosine receptor for olfaction in fish. Curr. Biol. 27, 1437–1447.e4 (2017).

29. Hussain, A. et al. High-affinity olfactory receptor for the death-associated odor cadaverine. Proc Natl Acad Sci USA 110, 19579–19584 (2013).

30. Diaz-Verdugo, C., Sun, G. J., Fawcett, C. H., Zhu, P. & Fishman, M. C. Mating suppresses alarm response in zebrafish. Curr. Biol. 29, 2541–2546.e3 (2019).

31. Yabuki, Y. et al. Olfactory receptor for prostaglandin F2α mediates male fish courtship behavior. Nat. Neurosci. 19, 897–904 (2016).

32. Braubach, O. R., Wood, H.-D., Gadbois, S., Fine, A. & Croll, R. P. Olfactory conditioning in the zebrafish (Danio rerio). Behav. Brain Res. 198, 190–198 (2009).

33. Frank, T., Mönig, N. R., Satou, C., Higashijima, S.-I. & Friedrich, R. W. Associative conditioning remaps odor representations and modifies inhibition in a higher olfactory brain area. Nat. Neurosci. 22, 1844–1856 (2019).

34. Herrera, K. J., Panier, T., Guggiana-Nilo, D. & Engert, F. Larval zebrafish use olfactory detection of sodium and chloride to avoid salt water. Curr. Biol. 31, 782–793.e3 (2021).

35. Krishnan, S. et al. The right dorsal habenula limits attraction to an odor in zebrafish. Curr. Biol. 24, 1167–1175 (2014).

36. Doyle, W. I. & Meeks, J. P. Excreted steroids in vertebrate social communication. J. Neurosci. 38, 3377–3387 (2018).

37. Kadakia, N. et al. Odour motion sensing enhances navigation of complex plumes. Nature 611, 754–761 (2022).

38. Felsenberg, J., Barnstedt, O., Cognigni, P., Lin, S. & Waddell, S. Re-evaluation of learned information in Drosophila. Nature 544, 240–244 (2017).

39. Qiu, Q., Wu, Y., Ma, L. & Yu, C. R. Encoding innately recognized odors via a generalized population code. Curr. Biol. 31, 1813–1825.e4 (2021).

40. Iravani, B., Schaefer, M., Wilson, D. A., Arshamian, A. & Lundström, J. N. The human olfactory bulb processes odor valence representation and cues motor avoidance behavior. Proc Natl Acad Sci USA 118, (2021).

41. Mathuru, A. S. et al. Chondroitin fragments are odorants that trigger fear behavior in fish. Curr. Biol. 22, 538–544 (2012).

42. Masuda, M. et al. Identification of olfactory alarm substances in zebrafish. Curr. Biol. 34, 1377–1389.e7 (2024).

43. Yilmaz Balban, M., et al. Human responses to visually evoked threat. Curr. Biol. 31, 601–612.e3 (2021).

44. Kuroda, S., Lalonde, R. L., Mansour, T. A., Mosimann, C. & Nakamura, T. Multiple embryonic sources converge to form the pectoral girdle skeleton in zebrafish. Nat. Commun. 15, 6313 (2024).

45. Moyle, P. & Jr, J. C. Fishes: An Introduction to Ichthyology. 752 (Pearson, 2003).

46. Ahrens, M. B., Orger, M. B., Robson, D. N., Li, J. M. & Keller, P. J. Whole-brain functional imaging at cellular resolution using light-sheet microscopy. Nat. Methods 10, 413–420 (2013).

47. Baden, T. From water to land: Evolution of photoreceptor circuits for vision in air. PLoS Biol. 22, e3002422 (2024).

48. Nady, A., Peimani, A. R., Zoidl, G. & Rezai, P. A microfluidic device for partial immobilization, chemical exposure and behavioural screening of zebrafish larvae. Lab Chip 17, 4048–4058 (2017).

49. Mattern, K., von Trotha, J. W., Erfle, P., Köster, R. W. & Dietzel, A. NeuroExaminer: an all-glass microfluidic device for whole-brain in vivo imaging in zebrafish. *Commun*. Biol. 3, 311 (2020).

50. Candelier, R. et al. A microfluidic device to study neuronal and motor responses to acute chemical stimuli in zebrafish. Sci. Rep. 5, 12196 (2015).

51. van Giesen, L., Neagu-Maier, G. L., Kwon, J. Y. & Sprecher, S. G. A microfluidics-based method for measuring neuronal activity in Drosophila chemosensory neurons. Nat. Protoc. 11, 2389–2400 (2016).

52. Friedrich, R. W. & Korsching, S. I. Combinatorial and chemotopic odorant coding in the zebrafish olfactory bulb visualized by optical imaging. Neuron 18, 737–752 (1997).

53. Kim, D. H. et al. Pan-neuronal calcium imaging with cellular resolution in freely swimming zebrafish. Nat. Methods 14, 1107–1114 (2017).

54. Flash, T. & Hogan, N. The coordination of arm movements: an experimentally confirmed mathematical model. J. Neurosci. 5, 1688–1703 (1985).

55. Todorov, E. & Jordan, M. I. Smoothness maximization along a predefined path accurately predicts the speed profiles of complex arm movements. J. Neurophysiol. 80, 696–714 (1998).

56. Anticipatory behavior in adaptive learning systems: foundations, theories, and systems. vol. 2684 (Springer Berlin Heidelberg, 2003).

57. Dunn, T. W. et al. Brain-wide mapping of neural activity controlling zebrafish exploratory locomotion. eLife 5, e12741 (2016).

58. Horstick, E. J., Bayleyen, Y. & Burgess, H. A. Molecular and cellular determinants of motor asymmetry in zebrafish. Nat. Commun. 11, 1170 (2020).

59. Gillis, G. B. Undulatory Locomotion in Elongate Aquatic Vertebrates: Anguilliform Swimming since Sir James Gray. Am. Zool. 36, 656–665 (1996).

60. Haesemeyer, M., Robson, D. N., Li, J. M., Schier, A. F. & Engert, F. The structure and timescales of heat perception in larval zebrafish. Cell Syst. 1, 338–348 (2015).

61. Balakrishnan, K. A. & Haesemeyer, M. Behavioral and circuit principles of temperature gradient navigation. Curr. Biol. 35, 5395–5410.e8 (2025).

62. Carbo-Tano, M. et al. The mesencephalic locomotor region recruits V2a reticulospinal neurons to drive forward locomotion in larval zebrafish. Nat. Neurosci. 26, 1775–1790 (2023).

63. Berg, E. M. et al. Brainstem circuits encoding start, speed, and duration of swimming in adult zebrafish. Neuron 111, 372–386.e4 (2023).

64. Mueller, T., Dong, Z., Berberoglu, M. A. & Guo, S. The dorsal pallium in zebrafish, Danio rerio (Cyprinidae, Teleostei). Brain Res. 1381, 95–105 (2011).

65. Kermen, F., Franco, L. M., Wyatt, C. & Yaksi, E. Neural circuits mediating olfactory-driven behavior in fish. Front. Neural Circuits 7, 62 (2013).

66. Korsching, S. I. Taste and smell in zebrafish. in The senses: A comprehensive reference 466–492 (Elsevier, 2020). doi:10.1016/B978-0-12-809324-5.24155-2.

67. Thelen, E. Rhythmical behavior in infancy: An ethological perspective. Dev. Psychol. 17, 237–257 (1981).

68. Wasserman, S. M. et al. Olfactory neuromodulation of motion vision circuitry in Drosophila. Curr. Biol. 25, 467–472 (2015).

69. Frye, M. A. & Dickinson, M. H. Motor output reflects the linear superposition of visual and olfactory inputs in Drosophila. J. Exp. Biol. 207, 123–131 (2004).

70. Mayseless, O., et al. Odor preference maps to cohesive transcriptional domains in the olfactory bulb. BioRxiv (2026) doi:10.64898/2026.06.11.731560.

71. Jenkins, B. & Frank, T. Distributed representations of chemosensory valence in a naïve vertebrate brain. BioRxiv (2026) doi:10.64898/2026.06.17.732810.

72. Leung, L. C. et al. Neural signatures of sleep in zebrafish. Nature 571, 198–204 (2019).

73. Yang, C. et al. A population code for spatial representation in the zebrafish telencephalon. Nature 634, 397–406 (2024).

74. Fotowat, H., Lee, C., Jun, J. J. & Maler, L. Neural activity in a hippocampus-like region of the teleost pallium is associated with active sensing and navigation. eLife 8, (2019).

75. Trinh, A.-T. et al. Hierarchical sensory processing in zebrafish thalamocortical-like circuits. Science eaec2171 (2026) doi:10.1126/science.aec2171.

76. Lifshitz, I. et al. Distinct distributed neural dynamics predict pallium-dependent social approach. Nat. Commun. 17, (2026).

77. Smith, C. D. et al. Negative-Valence Neurons in the Larval Zebrafish Pallium. eLife (2026) doi:10.7554/eLife.110720.1.

78. Brazeau, M. D. & Friedman, M. The origin and early phylogenetic history of jawed vertebrates. Nature 520, 490–497 (2015).

79. Veith, J. et al. The mechanism for directional hearing in fish. Nature 631, 118–124 (2024).

80. Fehrenbach, J., Weiss, P. & Lorenzo, C. Variational algorithms to remove stationary noise: applications to microscopy imaging. IEEE Trans. Image Process. 21, 4420–4430 (2012).

81. Pnevmatikakis, E. A. & Giovannucci, A. NoRMCorre: An online algorithm for piecewise rigid motion correction of calcium imaging data. J. Neurosci. Methods 291, 83–94 (2017).

82. Giovannucci, A. et al. CaImAn an open source tool for scalable calcium imaging data analysis. eLife 8, (2019).

83. Hernandez, R. E., Galitan, L., Cameron, J., Goodwin, N. & Ramakrishnan, L. Delay of initial feeding of zebrafish larvae until 8 days postfertilization has no impact on survival or growth through the juvenile stage. Zebrafish 15, 515–518 (2018).

84. Christou, M., Kavaliauskis, A., Ropstad, E. & Fraser, T. W. K. DMSO effects larval zebrafish (Danio rerio) behavior, with additive and interaction effects when combined with positive controls. Sci. Total Environ. 709, 134490 (2020).

85. Rhodes, M. Introduction to Particle Technology. (1989).

86. Randlett, O. et al. Whole-brain activity mapping onto a zebrafish brain atlas. Nat. Methods 12, 1039–1046 (2015).

87. Ramirez, A. D. & Aksay, E. R. F. Ramp-to-threshold dynamics in a hindbrain population controls the timing of spontaneous saccades. Nat. Commun. 12, 4145 (2021).

88. Moon, Y. I., Rajagopalan, B. & Lall, U. Estimation of mutual information using kernel density estimators. Phys. Rev. E Stat. Phys. Plasmas Fluids Relat. Interdiscip. Topics 52, 2318–2321 (1995).

